# SpectroVQ: Noise-Aware Compression of Proteomics Data via Vector-Quantized Deep Learning improves MS/MS data storage and Peptide Identification

**DOI:** 10.64898/2026.09.05.749393

**Authors:** James H.W. Li, Ayman Hoque, Henry Lam

**Author notes:** Correspondence to: Prof. Henry Lam.

## Abstract

The amount of proteomics data generated has dramatically grown for the past decade due to the wider accessibility to mass spectrometers and technological advances. Current data storage and compression techniques largely treat mass spectra as meaningless series of numbers, wasting storage on useless noise and limiting the compression ratio. Here, we present SpectroVQ, a noise-aware vector-quantized autoencoder to compress and denoise peptide tandem mass spectra without any prior annotation by exploiting peptide fragmentation pattern using deep-learning Evaluation results showed that SpectroVQ can preferentially retain useful signals in spectra from diverse peptide ions, including those in unseen datasets. SpectroVQ achieved over 3-fold increase in compression ratio over mzMLb while maintaining over 0.9 in average cosine similarity and 90% agreement in peptide identifications. In addition, we develop a novel strategy to increase peptide identifications by ∼15% via ordinary library searching, by leveraging the tuneable denoising capability of SpectroVQ.

## Introduction

Liquid chromatography (LC) coupled to mass spectrometry (MS) has emerged as one of the most popular platform for proteomics^1^. Thanks to rapid technological advances and the wider adoption of proteomics in life science research, the amount of MS data generated has grown exponentially in the past two decades. To facilitate open science practices, massive amount of mass spectrometry proteomics data are being uploaded to public data repositories under the ProteomeXchange consortium^2^ (Figure 1a), with the PRIDE repository (https://www.ebi.ac.uk/pride/) receiving more than 6000 data submission in 2025 alone^3^. It is expected that the trend will continue in the foreseeable future. Therefore, efficient storage for proteomics data is an increasingly pressing need.

**Figure 1:**
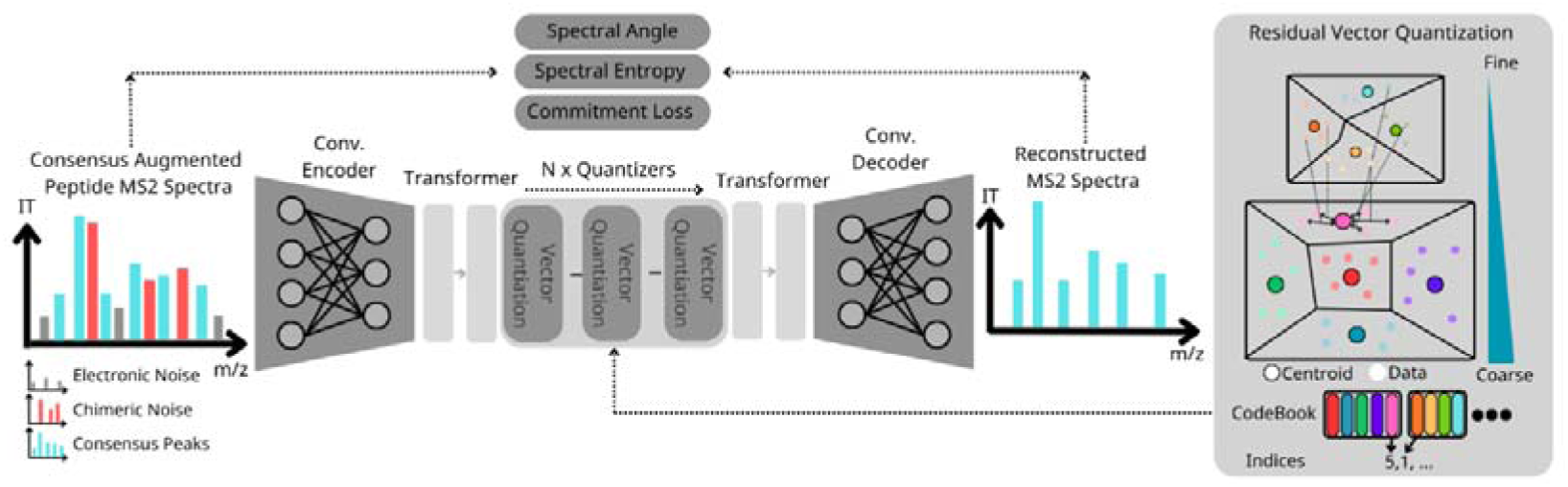
Overview of SpectroVQ. SpectroVQ consists of three main parts, the encoder, residual vector quantizers (RVQs) containing multiple quantizers, each with a separate codebook, and a decoder. The encoder and decoder consist of a combination of convolution layers and transformer Layers. In the RVQs, the vector is first quantized to its nearest centroid in the codebook. The difference between the centroid and vector is then quantized, and the whole process repeats for N times, forming a series of quantizers. Centroid vectors are stored internally as the codebook for SpectroVQ and are constructed during training, in which SpectroVQ is trained to remove chimeric and electronic noises in augmented noisy spectra. During the inference stage, experimental spectrum is first passed through the encoder to extract noise-free features in the latent space and then undergo quantization with the RVQs. The indices of the quantized latents used in the codebook are stored in the compressed data. To decompress the spectrum, the indices are re-projected to the codebook and pass through the decoder. Since ideally only the de-noised features are stored in the latent vector, the reconstructed spectrum would not contain noise peaks.

Current standards in mass spectrometry storage rely on representing mass spectra as a sequence of peaks, each encoded by a pair of numbers, the mass-to-charge ratio (m/z) and the intensity. The open standardized format mzML, developed under the Proteomics Standards Initiative (PSI)^4^ in the Human Proteome Organization (HUPO), laid the foundation for vendor-independent mass spectrometry data storage. In mzML, the metadata of each MS scan is stored as text in the XML format under a strict standard vocabulary in the form of cvParam. The spectrum itself is stored as two binary base 64 data arrays, one representing m/z values and the other representing intensity values^4^. As data volume grows, the plain text-based storage mechanism of mzML has become increasingly burdensome. In response, various data standards and compression methods were developed over the past decade. Whereas the metadata can easily be stored more efficiently in binary representation or relational databases than in XML format, better ways to store the MS spectra, which usually take up the vast majority of a typical mzML file, would be more impactful. To that end, compression methods such as MassComp^5^ perform lossless compression by numerical transforming m/z and intensity values using delta and arithmetic encoding, whereas later methods MSpack^6^ and mzMLb^7^ performs lossy compression by reducing the numerical precision in m/z and intensity values. Alternatively, mzPeak^8^ unifies multiple compression techniques by allowing both lossless (point layout) and lossy (chunked layout) compression.

Although these techniques achieved impressive compression performance, they are inherently treating mass spectra as a series of numbers, without any knowledge of the fragmentation patterns of peptides. Fundamentally, the primary use of a peptide tandem mass spectrum is peptide identification, and not all peaks in a spectrum is equally useful for that purpose. Interleaved with “signal” peaks that enable peptide identification are a greater number of “noise” peaks arising from instrument noise or chemical contaminants in the sample^9^. Storing these extra “noise” peaks in each spectrum unnecessarily increases the burden on data repositories. Moreover, removing these noise peaks can be beneficial for downstream data analysis in both proteomics^10,11^ and metabolomics^12,13^. Therefore, an ideal compression method should prioritize peaks by utility in a hierarchical structure, letting use trade compression ratio for detail. Crucially, this hierarchical compression method also functions as a denoising tool, potentially improving downstream analysis even when storage is not the primary concern.

Therefore, in this work, we introduced SpectroVQ, the first-of-its-kind deep learning model for compressing and denoising peptide tandem mass spectra. Unlike existing methods for data compression, which are limited to redundancy reduction and numerical transformations, SpectroVQ focuses on maximizing semantic information retention, rather than on strict reconstruction fidelity. Inspired by recent advances in deep learning-based audio compression methods called “codecs,” which seek to distill the human-perceptible sounds in soundtracks rather than transforming the original sound waves. SpectroVQ attempts to do the same to mass spectra. In brief, it encodes the mass spectra in a dense embedding space and subjects the embeddings to a series of vector quantization steps. This enable SpectroVQ to efficiently converts a spectrum into a series of integers, which can be decompressed to a reconstructed mass spectrum that is highly similar to the original spectrum. As a result, far fewer bits are needed to store spectra than using binary or floating numbers to directly represent the m/z and intensity values. In addition, SpectroVQ was trained with a tuneable denoising capability. Users can reconstruct multiple versions of the same spectrum by specifying the number of quantizers used, which provides flexibility in storage sizes and improves peptide identification rates.

## Results

### SpectroVQ Overview

The architecture of SpectroVQ consists of a convolution neural network (CNN) and a transformer block as the encoder, a corresponding symmetrical decoder, and a series of residual vector quantizers (RVQs) in between^14^. SpectroVQ takes a binned MS2 spectrum as its input, encodes the whole spectrum as a lower-dimension embedding vector, a representation of the spectrum that preferentially retains the signals via the CNN. The embedding is then fed to the series of RVQs. Each RVQ comprises a codebook, which contains the centroids of all embeddings seen during training. For each spectrum to be compressed, the embedding is mapped to, and approximated, by its geometrically nearest vector contained in each codebook, whose integer index is stored. Then, the difference between the vector and its projection – the residual vector – passes through the next quantizer in the sequence, where the information contained gradually decreases, forming a hierarchy (Figure 1). Finally, the integer indices of the closest codebook entry of all the quantizers are stored as “tokens” for that spectrum, and are all that are required for reconstruction.

To encourage SpectroVQ to retain signals preferentially, we employ a noise-to-clean training strategy (Supplementary Figure S1). We first augment high-quality consensus library spectra generated by SpectraST^15^ with simulated electronic noise (due to inherent random fluctuations of electrical signals detected by the mass spectrometer) and contaminant peaks from co-fragmenting peptides, then task the model to reconstruct the original unaugmented consensus spectra (Figure 1c). It is assumed that the unaugmented consensus spectra contain mostly “signals” worth retaining. With this training goal, the encoder and the codebook of the RVQs will be updated progressively to capture more “signals” while discarding noise. The peptide identifications of the spectra are hidden from the model and play no role in SpectroVQ. Therefore, what the model learns are general rules about peptide fragmentation patterns and how to distinguish signals from noise irrespective of the peptide identification of the spectra.

One important design criterion for SpectroVQ is hierarchical data compression, meaning that the most informative features of the data are retained first. This is accomplished by dropping out the last-N quantizers during training at random. This challenges the model to retain as much information in the earlier quantizers as possible.

To evaluate the success of hierarchical compression, we looked at the reconstruction performance using different numbers of quantizers (Figure 2a). As expected, with more quantizers, the decoder receives more information from the latent embedding to reconstruct a spectrum with higher fidelity, at the expense of more storage space. However, the increment in reconstruction fidelity diminishes as more quantizers are used. Addition of the second quantizer results in a median spectral similarity increase of 0.11, whereas the inclusion of the last (12^th^) quantizer only improves median spectral similarity by 0.02 (Figure 2b). This demonstrated that the hierarchical compression works as intended. For subsequent evaluations of SpectroVQ, we choose to use 4 quantizers, corresponding to the storage size of 13.8 kilobytes per spectrum, which we believe strikes a good balance between reconstruction fidelity and storage size.

**Figure 2:**
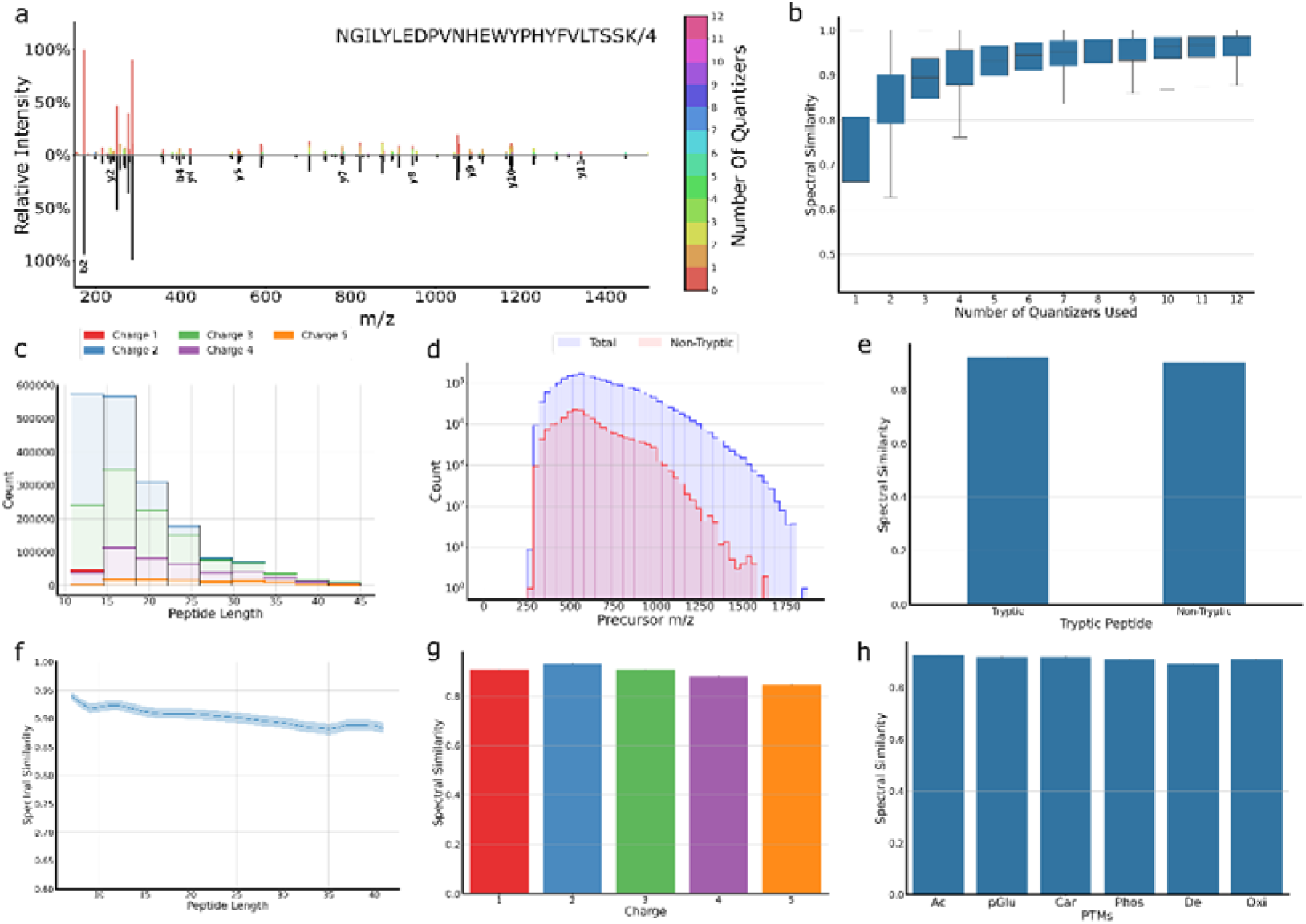
Reconstruction Performance using SpectroVQ. (**A)** Example reconstructed spectrum with SpectroVQ using multiple numbers of quantizers. The graph illustrates reconstructed spectrum using different number of quantizers. Increasing number of quantizers lead to an increased number of peaks retained. However, in this example, using the first 2 quantizers already cover the majority of b,y fragments in the spectrum, suggesting SpectroVQ’s ability to implicitly rank peaks. **(B)** Distribution of spectral similarity (defined as cosine similiarity based on square-root transformed intensities) between original consensus and reconstructed spectrum at different levels of quantization. Boxes indicate interquartile range. The first few quantizers (1-4) contain most of the information from the spectrum, yielding a higher gain in spectral similarity when they are included. **(C)** Distribution of spectra from peptides with different lengths and charge states in the consensus library for training the model. The majority of spectra are from peptide with precursor charges 2+ and 3+ and lengths below 20. **(D)** The proportions of tryptic and non-tryptic spectra in the consensus library by precursor m/z. **(E-H)** Average spectral similarity between the original consensus and reconstructed spectrum, broken down by trypticness **(E)**, peptide lengths **(F)**, charge states **(G)** and post-translational modifications **(H)** using 4 quantizers. As each spectrum is compressed to occupy the same storage size, a slightly drop in performance is observed in denser spectra with more informative peaks, such as those of longer peptides and higher charge states.. **Abbreviations**: Acetylation (Ac), Pyroglutamic acid modification (eGlu), Cysteine Carbamidomethylation (Car), Phosphorylation (Phos), Asparagine and Glutamine Deamidation (De), Methonine Oxidation (Ox)

### SpectroVQ Enables Consistent Reconstruction of Peptide Tandem Mass Spectra

We next assessed if SpectroVQ shows consistent reconstruction performance for all peptide tandem mass spectra, without bias towards peptides of different attributes, such as precursor mass, charge state, length and modification. Note that since our model is trained on higher-energy collisional dissociation (HCD) data acquired with data-dependent acquisition (DDA), we focused on evaluating this common type of peptide tandem mass spectra. We subjected human consensus library spectra from PeptideAtlas (https://db.systemsbiology.net/sbeams/cgi/PeptideAtlas/buildDetails?atlas_build_id=607) to SpectroVQ and measured the spectral similarity between the reconstructed spectrum and the original library spectrum. Overall, SpectroVQ achieves excellent reconstruction fidelity across the entire library. SpectroVQ performs well for tryptic and non-tryptic peptide spectra with average spectral similarity above 0.9 (Figure 2e).

Spectral similarity across different peptide length shows similar performance with values above 0.84, with only a slight drop in longer peptides. Furthermore, reconstruction fidelity across multiple post-translational modifications (PTMs) is consistently high. Evaluation on the testing portion also shows identical conclusions, thus showing that the good performance is not due to overfitting (Supplementary Figure S2).

### SpectroVQ Removes Unwanted Noise after Reconstruction

As part of our goal is to retain semantic information related to the underlying peptide only without preserving all original peaks, we seek to assess SpectroVQ’s denoising ability by comparing the portion of canonical ions (defined as b, y ions of charge states 1+ or 2+) between the original augmented spectrum and the corresponding reconstructed spectrum. An example augmented spectrum from the testing portion of the PeptideAtlas dataset (Figure 3a) illustrates how SpectroVQ removes most of the noisy peaks added while preserving the original b and y ion fragment ladder. Similar performance was also seen in the entire testing dataset. The average canonical ion fraction over the entire testing dataset, improves from 5% to 13%, while the average number of peaks drops from 244.4 to 87.4. (Figure 3b). Comparison with other naïve denoising strategies, such as thresholding the peaks at 1% of the maximum intensity or keeping only the 100 or 150 most intense peaks, also shows SpectroVQ’s better denoising performance (Figure 3c). Notably, we observed that SpectroVQ’s denoising ability outperform top N filters for different levels of augmented noise in the same testing dataset, where N is chosen to match the number of peaks in SpectroVQ’s reconstructed spectrum (Supplementary Figure S3).

**Figure 3:**
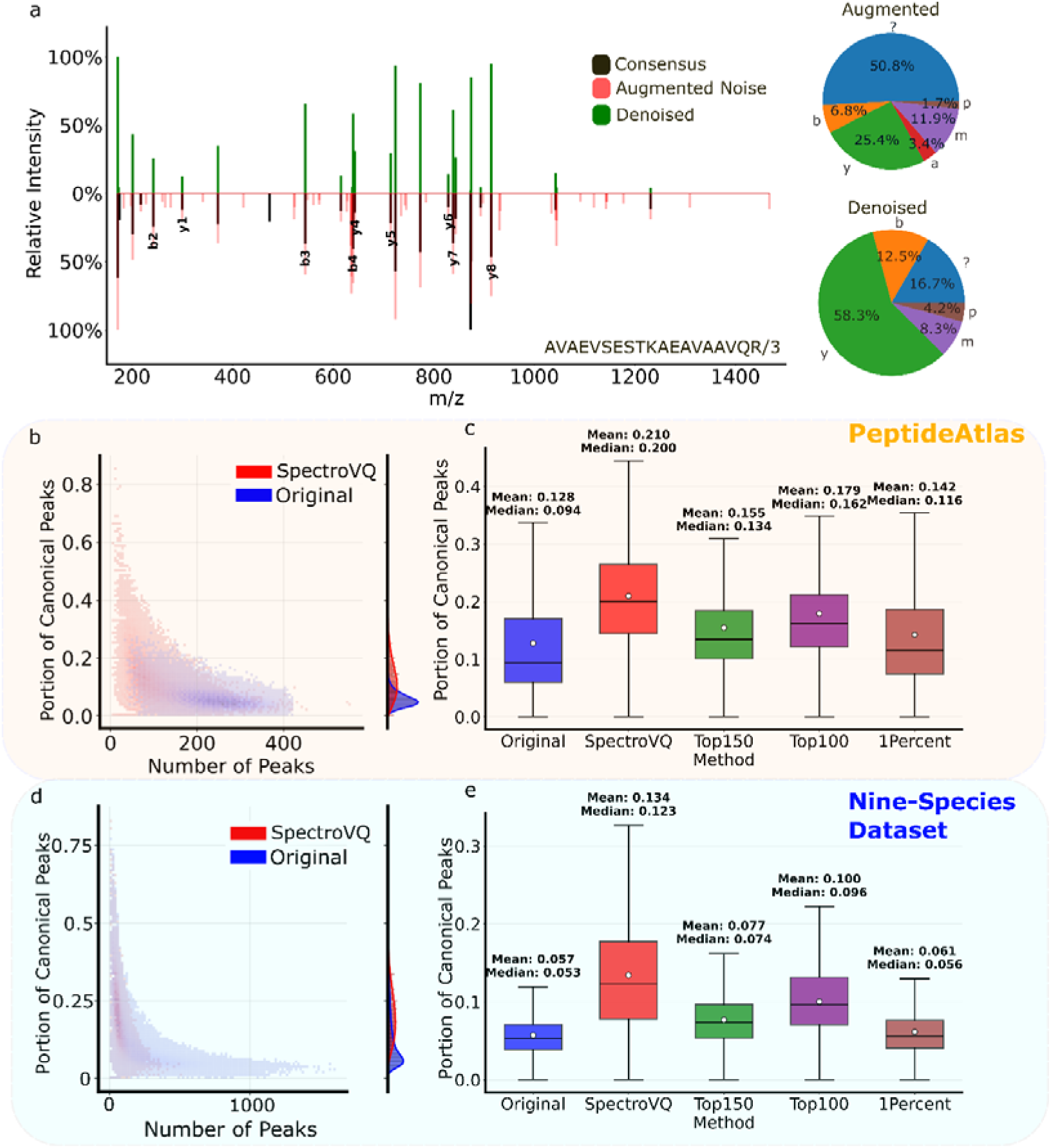
Denoising Performance of SpectroVQ. **(A)** An example of a denoised spectrum, plotted against the augmented spectrum with peaks from the original consensus spectrum (“SpectroVQ”) nd the added artificial noise (“Augmented Noise”). The distribution of annotated ion types of the augmented spectrum and the denoised spectrum are shown on the right. Added noise peaks in the augmented spectrum are largely removed by SpectroVQ during reconstruction while the signal peaks originally present in the consensus spectrum are retained. The increase in the fraction of annotated peaks (including b, y ions, precursor losses (p) and internal fragments (m)) after SpectroVQ reconstruction indicates SpectroVQ’s ability to retain sequence-specific information and reduce noise. **(B)** Fraction of canonical peaks vs. the number of peaks before and after SpectroVQ denoising of the PeptideAtlas dataset. **(C)** Fraction of canonical peaks in the original spectrum and after denoised by SpectroVQ or other strategies. **(D-E)** Corresponding plots for the nine-species dataset not used in training. SpectroVQ greatly reduces the number of peaks while boosting the fraction of canonical ion peaks and consistently outranks other baseline methods of picking the top 150 peaks (Top150), the top 100 peaks (Top100), and thresholding at 1% of the base peak intensity (1Percent) in both datasets.

We also evaluated SpectroVQ on data from other organisms, to demonstrate that the model is not overfitted to our training data, which is exclusively from human samples. On the nine-species dataset^16^, SpectroVQ shows similar trend with that in our human testing dataset, showing an average increase from 0.13 to 0.21 while the average number of peaks drop from 232 to 64 (Figure 3d). Again, SpectroVQ outperforms the other strategies, resulting in the highest canonical ion fraction overall (Figure 3e). SpectroVQ also have the highest canonical ion fraction in every species, showing it’s the model’s ability to generalize to other unseen datasets (Supplementary Figure S4 & Supplementary Figure S5).

### SpectroVQ Captures Peptide Fragmentation Patterns in the Quantized Latent Space

We further investigated whether SpectroVQ’s embeddings in the latent space carry sufficient information to distinguish spectra from slightly different peptides. Using another dataset for which replicate spectra from the same peptide ion are present (Method), we compressed them using SpectroVQ and performed principal component analysis (PCA) on the embeddings. Even through spectra embeddings are quantized, the resultant embedding still retain enough information to separate spectrum from different peptide. The PCA plot shows that spectra of the same peptide identification are clustered together, and a clear separation between spectra of different identifications can be observed (Figure 4a). Compared to a similar analysis using the original spectrum (vectorized with bin size 0.1 Th), SpectroVQ achieves better clustering performance in both the Silhouette score (p <0.001) and David-Bould score (p <0.001).

**Figure 4:**
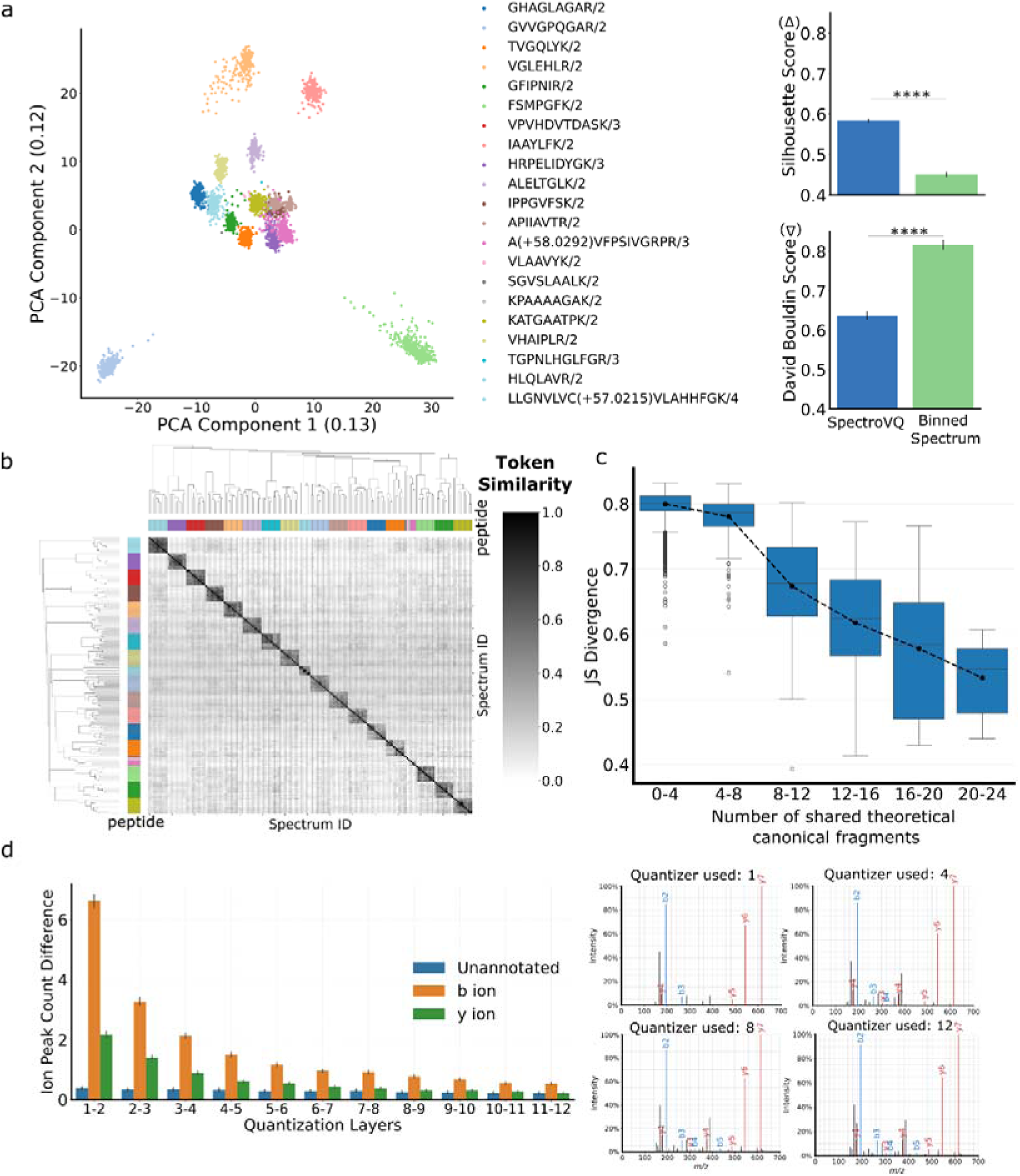
Peptide information contained in the latent space of SpectroVQ. **(A)** (left) PCA plot of the latent embeddings of selected spectraannotated with different peptides. (right) Comparisons of the Silhouette score (Top) and David-Bouldin score (Bottom) between clusters of SpectroVQ latent embeddings and those of the original binned spectra. Without knowledge of the peptide identifications, the latent embeddings already form distinct clusters by PCA using only two components. Improvements in Silhousette score and David-Bouldin score of the clustering shows that the latent embeddings are better able to capture peptide-specific information in the spectra than the raw spectra themselves(**** p-value < 0.001) **(B)** Heatmap of pairwise similarities of tokens (i.e., indices of closest entries of the codebook) of the first quantizer of the spectra included in (A). Token similarity is defined as the jaccard index between the tokens from two spectra. Spectra of the same peptide identification are highly similar in their tokens, whereas there is almost no overlap in the tokens between spectra of different peptides. (C) Boxplot of Token Distribution Divergence between spectra groups from different peptide against the theoretical number of shared canonical fragments. We compared the Jensen-Shannon divergence between tokens distributions from different groups of peptide from the Human Proteome Project dataset. Peptides which shares no theoretical fragments shows the highest divergence in their token distributions. With a higher number of shared canonical fragments, JS divergence between token distribution gradually minimizes. This further suggest that learnt latent token distribution correlates with canonical fragmentation pattern. **(D)** Differences in ion count (b,y, unannotated) across multiple quantization layerss. A decreasing trend in b,y ion is observed when a quantization layer is added in late series, this suggest that more important b,y fragments are ranked and stored in the earlier stages in the series of quantizers, creating a hierarchical storage.

We also observed that the quantized vector (namely, the “tokens” representing the closet codebook entries of all quantizerst) carries meaningful peptide information. We compared the non-conditioned tokens of the first quantizer representing replicates (spectra identified to the same peptide ion) included in Figure 4a. Even though peptide information is not passed into the model, replicate spectra share a high average token similarity (Same peptide: 0.503; Different peptides: 0.0371), explaining the clustering patterns in the PCA plot. Furthermore, distribution changes to these tokens can reveal the changes to the underlying peptide sequence. Spectra from peptides with no overlapping fragments yields a high divergence in their token distribution (Figure 3c). With more overlapping canonical fragments, we observed a gradually decreasing divergence between their token distribution (Figure 4c).

Finally, we observed that SpectroVQ learned to distinguish peptide fragmentation pattern not only in individual token but among the series of quantizers. Changes to the canonical ions are focused on earlier quantizers while the number of unannotated ions remains unchanged over the whole series of quantizers. This suggests that SpectroVQ implicitly ranked peaks during its compression, prioritizing the storage of canonical peaks at earlier RVQs and leaving other less crucial peaks at later RVQs. This is an indirect demonstration of how hierarchical data compression is achieved in SpectroVQ.

### Lossy Compression using SpectroVQ yields Similar Search Results

We evaluated SpectroVQ’s compression performance on an external real-life dataset from Bart et. al.^17^. This ensures SpectroVQ’s compression and denoising ability can be extended to real-life experimental datasets. SpectroVQ improves the compression size by over 4-folds compared to the compression method mzMLb^7^ (Figure 5a). In addition, despite sharing the same parquet file format, SpectroVQ also outperforms the other lossless format ArcMS^18^, also with more than 4-folds compression. Compared to the original file size in mzML, SpectroVQ achieves over 30-fold compression, with over 120-fold compression for some spectra with a large number of peaks. More importantly, SpectroVQ outperforms the baseline strategy of picking the top N peaks, with N chosen to match SpectroVQ’s compression ratio for that spectrum, preserving a higher average similarity between the original spectrum and the corresponding reconstructed spectrum. (SpectroVQ:0.88, TopN:0.64). (Figure 5b).

**Figure 5:**
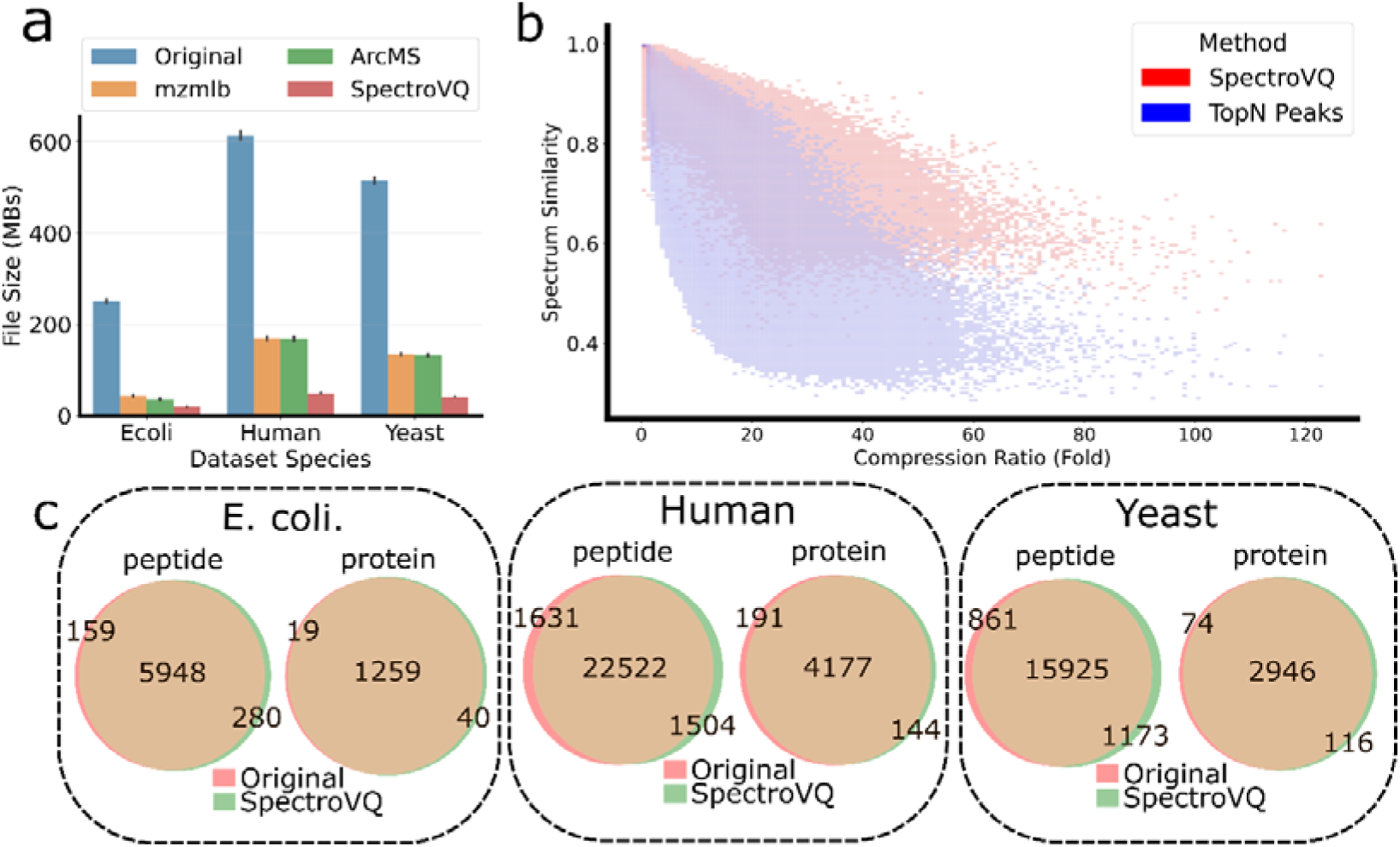
Testing SpectroVQ’s compression on the HYE external dataset. **(A)** Barplots showing file sizes of experimental data in mzML, mzMLb, ArcMS and SpectroVQ. SpectroVQ demonstrate excellent compression ability, which reducees the file sizes over 5-folds compared to mzMLb and ArcMS. **(B)** Spectral similarity between the original and denoised spectra vs. the compression ratio of individual spectra, comparing SpectroVQ versus the baseline strategy of picking top N peaks, with N chosen to match the same compression ratio achieved by SpectroVQ for that spectrum. SpectroVQ is able to condense more information in the same amount of storage space, leading to a higher spectral similarity with the original spectrum. **(C)** Venn diagram of unique peptides and proteins identified in the *E. coli*. (Top), Human (Middle) and Yeast (Bottom) portions of the three-species dataset, showing the clear overlap of identifications between the original data and the SpectroVQ-compressed and reconstructed data. This demonstrates that minimal biological information is lost during the compression by SpectroVQ.

We then compared the peptide identification results of SpectroVQ’s compressed and reconstructed data against those of the original data. Although mzMLb compressed lossily, the loss in m/z numerical precision in mzMLb^7^-compressed data did not alter the search results. For SpectroVQ, a high agreement was observed in both the peptide and protein level among confident identifications filtered at 1% false discovery rate (FDR) at the respective levels) (Figure 5c). Over 90% of the identified peptide and proteins are shared between the search results, with a slight gain in identifications for the *E. coli* and Yeast datasets and only a slight loss for the human dataset after SpectroVQ reconstruction. This similarity is also observed at the spectrum level (Supplementary Figures S6)

### Novel Denoising Strategy Significantly Improve Search Results

In our previous tests, we used 4 residual vector quantizers as the optimal number to perform compression, striking a balance between reconstruction fidelity and compression ratio. Presumably, varying the number of quantizers will lead to different degrees of denoising and different peaks being retained after compression and reconstruction, which will in turn affect whether marginal peptide identifications will be declared positive. Therefore, if data compression is not the sole objective, SpectroVQ’s tunable denoising ability can be leveraged to maximize peptide identifications for the same dataset (Figure 6a). Given an experimental spectrum, we produce 12 different versions of it with different levels of denoising by processing it with SpectroVQ using different number of quantizers, and search all of them. This is somewhat analogous to deploying multiple search engines or using different search parameters on the same dataset and combine the results.

**Figure 6:**
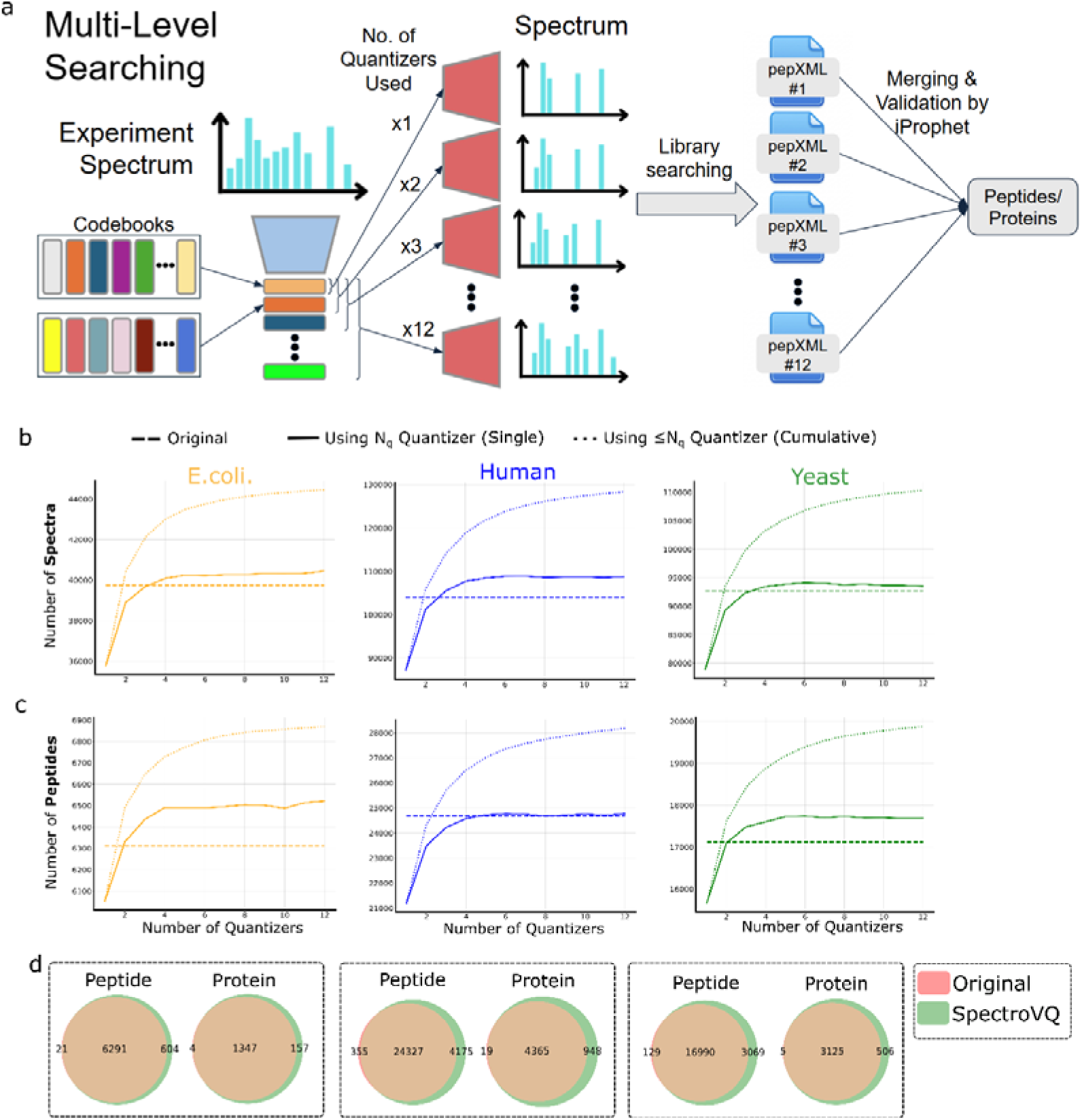
Library Searching using novel multi-level denoising strategy with SpectroVQ. **(A)** Multi-level denoising via varying the number of quantizers used in SpectroVQ. Instead of using an optimal number of quantizers, this strategy creates 12 versions of the same spectrum with different levels of denoising by varying the number of quantizers, and combines the search results of all the versions by InterProphet(B-C) The number of peptide-spectrum matches (B) and unique peptides (C) identified at 0.01 false-discovery rate (FDR) obtained from spectra at different denoising levels (1 = using first quantizer, 2 = using first two quantizers, etc.), showing numbers at each level (dotted line), and cumulatively (filled line). The identification number from only searching the original data is shown as a dashed line for comparison. Compared to the baseline of searching with the original data, the number of unique peptide and spectrum generally increases with higher number of quantizers used, reaching a plateau at about 4 quantizers. Cumulatively adding the identifications together at different denoising levels produces a substantial increase in the total number of spectra and peptide identified compared to the baseline. **(D)** Venn diagram of unique peptides and proteins identified in the E.coli. (Left), Human (Middle) and Yeast (Right) portion of the three-species dataset, showing the a consistent gain in peptide and protein idenfications using this strategy compared to the baseline.

We first compare the number of identifications using different number of quantizers. With higher number of quantizers, higher identification rates (at 1% FDR) are observed in all samples (Figure 6b). When we cumulatively add up identifications from spectra reconstructed using different number of quantizers, the identification rates show an increasing trend, suggesting that new unique peptide identifications are added from spectra at each level of denoising. Therefore, we used InterProphet^19^ to combine the search results intelligently under its Bayesian inference framework while controlling its FDR on the combined search results. This results in an increase in both the number of peptide identifications and the number of protein identifications in all 3 datasets, compared to the original data (Figure 6d). The number of spectra confidently identified at 1% FDR also increased more than 10% (Supplementary Figure S7). Closer inspection revealed that denoised spectra from all 12 levels of denoising (achieved by using 1 to 12 quantizers respectively) contributed to this improvement.

## Discussion

In this work, we introduce SpectroVQ, a deep learning-based autoencoder for compressing and denoising peptide tandem mass spectra. Our experiments suggest that SpectroVQ achieves the best compression ratio to date for proteomics data while maintaining excellent reconstruction fidelity, including for datasets not used in the training and for spectra of unseen peptides. Nearly identical search results can be obtained from the SpectroVQ-compressed and reconstructed data compared to the original data, while the storage needs are reduced more than 30-fold from mzML-formatted MS spectra employing base64 encoding. We also show that SpectroVQ learns how to distinguish signal from noise in typical peptide tandem mass spectra and can capture such information in its latent space, without knowing the peptide identification. The residual vector quantizers were trained to enable hierarchical data compression. We finally showed that SpectroVQ’s tuneable denoising capability can be leveraged to increase peptide and protein identifications, and in theory can be applicable to any identification pipeline

Unlike existing data compression methods in proteomics that treat mass spectra as series of meaningless numbers, SpectroVQ demonstrates great promise of lossy compression strategies that can take advantage of the inherent patterns of peptide fragmentation by deep learning. By training the model to distinguish signal from noise, one can achieve much better compression ratios while selectively retain useful information.

There are several directions to further improve on SpectroVQ. First, due to the model architecture, a binned spectrum corresponding to a fixed-length vector is required as the input (Methods). Although the m/z precision implied by our binning schemes already matches or exceeds that of most other deep learning models for MS data^20–24^, the loss in m/z precision would be detrimental to database searching performance, which is more sensitive to m/z values rather than the combination of intensity and m/z^25^ (Supplementary Figure S8). Second, SpectroVQ is currently trained on high-quality consensus library spectra from the human PeptideAtlas, which are exclusively compiled from HCD data acquired via data-dependent acquisition. Without further training, this version of SpectroVQ also achieves high reconstruction fidelity for TripleTOF 6600 DDA data, likely due to similar peptide fragmentation patterns (Supplementary Figure S9). For other data types, we expect that SpectroVQ can be trained to handle them, if sufficient high-quality training data is available, perhaps opening the possibility of compressing MS1 and data-independent acquisition (DIA) data as well. Third, our method focuses on the compressing the MS2 spectrum alone, without using other available information such as precursor mass, charge state, retention time and the corresponding MS1 spectrum. It is possible that these additional features can be leveraged to improve data compression and denoising performance.

Our work also opens new possibilities in data analysis in proteomics. First, SpectroVQ can act as the base model or feature extractors for other machine learning tasks in proteomics, such as spectrum prediction, post-translational modification detection and de novo sequencing. Unlike existing end-to-end training methods, incorporating SpectroVQ’s dedicated model for denoising can help improve the data quality for training and inference. Better time and space efficiency may also be attainable as the embeddings of SpectroVQ are shown to capture the essential information of the spectra in a much denser representation. Second, the token-based embeddings of SpectroVQ can potentially be used to generate realistic and artificial peptide tandem mass spectra for applications such as decoy generation and evaluation of search engines. As the fragmentation patterns of peptides are captured in the quantized latent space of SpectroVQ, resampling using an auxiliary model can in theory produce realistic spectra that follow the same patterns in real spectra. These tokens could potentially be used to represent MS data in large language models (LLMs). Instead of treating each peak in a spectrum as a token, using SpectroVQ’s rich spectrum representation can in theory improve LLMs’s understanding of MS data. Similar approaches of using alternative tokenization based on vector quantization rather than conventional text tokenization has been proven effective in many multimodal LLMs^26,27^. If the same trend applies to MS data, SpectroVQ can be a boost to current efforts to leverage LLMs and foundation models for proteomics data analysis.

## Methods

### Representation of the MS^2^ spectrum

Given a raw spectrum S, we encode the spectrum into as a one-dimensional vector by binning the spectrum with a binned width of 0.1 Th. We only included the peaks in the range between 150-1500 Th because peaks below 150 Th are not informative for peptide identification whereas few peaks exist beyond 1500 Th. Most database^25,28^ and library search engine^15^ uses peaks inside this range. These out-of-range peaks are still stored in the metadata file for reconstruction. The intensity of each spectrum is normalized, then their intensities are square-root transformed. This encourages the model to learn peaks with low intensities, similar to the approach in S. Goldman et.al.^29^.

### The architecture of SpectroVQ

The neural network architecture of SpectroVQ (Figure 1c) can be decomposed into three main modules, convolutional encoder, a residual vector quantization (RVQ) module and a convolutional decoder.

The architecture of the encoder and decoder is adopted from Encodec, a data compression model for soundtracks ^30^. The symmetrical encoder-decoder stack consisting of multiple residual convolutional blocks from SEANET^31^. Each convolutional block in SpectroVQ comprises a single residual 1D convolution block (2 convolutions of kernel size 8 with a skip-connection) and a strided convolution layer for down-sampling in the encoder or a transposed convolution for up-sampling in the decoder. Each up-sampling or down-sampling results apply a factor of 2 to the number of channels. All parameters in the convolutional block use weight normalization, and ELU is chosen as the activation function. Transformer blocks from Llama2 were used in the model to enhance sequential learning abilities, as suggested by its adoption in Mimi^32^. Each transformer block follows the typical structure of a self-attention layer followed by a feed-forward layer. Rotary embedding is used, following the example of Llama2, due to its excellent ability to capture long-range sequential information^33^. A final convolution layer is added before the RVQ module to further reduce the dimension of the compressed vectors. Ablation study on the transformer layers suggests this combination of convolution and transformer layers enhances both short- and long-range learning capabilities, in terms of capturing information in the binned spectrum (Supplementary Figure S10).

The latent vector outputted from the encoder is passed to the RVQ module. The RVQ module allows the latent vector to be projected to the closest vector in a learned codebook in a hierarchical manner. A total of 12 quantizers in series each with a codebook size of 1024 are present in SpectroVQ, though the actual number of quantizers used can be user-specified. In each quantizer, the latent embedding is replaced by the closest entry in the codebook. The residual (the difference between the latent embedding and the codebook entry) is then further quantized in the subsequent vector quantization layer (Algorithm 1) in a similar manner. To guide the model to learn a latent distribution that are more easily quantizable, mean-squared loss between the vector and the projections in the codebook (MSE) is used as the commitment loss as part of the loss function, which can guide the convolution autoencoders to generate vectors that are geometrically closer together given the same underlying peptide. The loss function will be further explained in section Loss function.

#### Algorithm 1

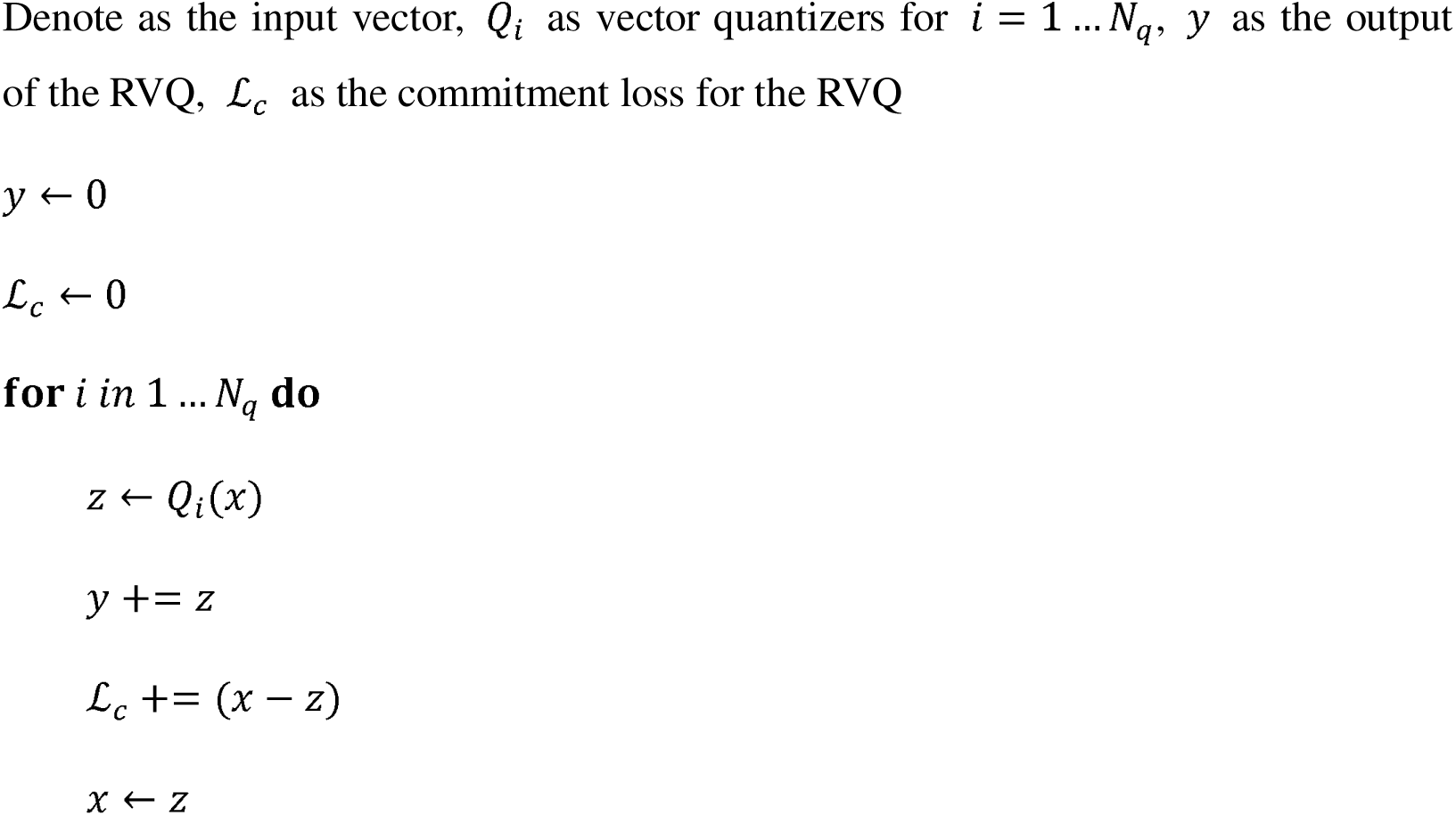

### Hyperparameter tuning

All hyperparameters in SpectroVQ are tuned using the Bayesian optimization framework Optuna^34^There are 2 objectives to be optimized, spectral similarity between reconstructed spectrum and original consensus spectrum, and quantized embedding length, which is roughly the number of indices to be stored in the final model. A longer embedding length often allows a higher spectral similarity as information is less compressed, but since our goal is to compress MS^2^ spectra, we want to optimize both contrasting objectives. Therefore, we try to locate the Pareto front and find the optimal hyperparameter to satisfy both goals. To facilitate the tuning process, we randomly sample 1% of spectra from the training dataset. The Pareto front plot is shown in Supplementary Figure S11. We choose the hyperparameter at elbow of the Parent front curve. The full hyperparameter list is available at Supplementary Table S1.

### Training Process

SpectroVQ is trained with a noise-to-clean approach (Supplementary Figure S1). Given a clean spectrum from the consensus library, we apply augmentation to simulate both electronic noises and peptide noises. Electronic noises are simulated by randomly generating peaks with intensities sampled from a Poisson distribution (*λ* = 4), following Kong et. al. ^35^. Peptide noises are simulated by adding another consensus spectrum randomly sampled from peptide with precursor mass range ±5Th of the original peptide. To ensure that the other spectrum does not overpower the original spectrum, we attenuate the second spectrum to a maximum of 0.4. Each augmentation has a 50% percent chance of applying to the original spectrum.

The codebook of the RVQ is simultaneously trained with the autoencoder during training. Following the approach from Defossez et. al.^30^, codebook entries are first initialized through a k-means clustering using the first training batch. In later batches, the entries are updated using an exponential moving average with a decay rate of 0.99. Straight-through estimator, or an identity function, is used to compute the gradient of the encoder during the backward phase. The number of quantizers used at each batch is sampled from a uniform distribution to form the hierarchy of the RVQs. The whole RVQ also have a 50% probability to be dropped out to stabilize training, as suggested by Defossez et. al.^32^.

The model is trained with the AdamW optimizer^36^. The initial learning rate is set to 1 x 10^−5^ The maximum number of epochs is set to 50. A warmup period of 5 epoch is set, followed with a cosine decay in learning rate until early stopping is triggered or the maximum number of epochs is reached. This warmup has been proven essential to training transformer-based models^37^. The batch size used was 32. A single Nvidia RTX4090 was used to train SpectroVQ, while another GPU Nvidia RTX3080Ti, along with the original Nvidia RTX4090, was utilized for inference.

### Loss function

The loss function includes the 3 terms, spectral angle *L_sa_*, spectral entropy*L_se_* and commitment loss *L_c_* Both spectral angle and spectral entropy^38^ accounts for the spectral quality while the commitment loss focuses on constructing a quantizable latent space. Formally, the loss can be written as:

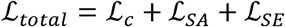

Spectral angle and spectral entropy loss are represented by the following equations:

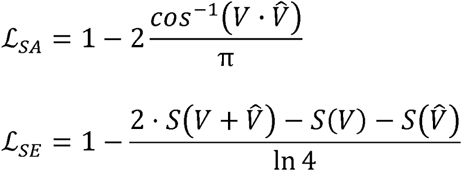

where:

*S(V) =-∑_v_ I_v_ ln(I_v_)* which *I_v_* denotes the intensity at each bin in the spectrum

### Compression with SpectroVQ

Experimental mzML files are parsed using Pyteomics^39^. Each MS2 spectrum read from the file is processed through the autoencoder of SpectroVQ to generate its latent embedding. The latent embedding is passed through the RVQ and the residual at each layer is matched to the trained codebook, and the closest entry is retrieved. Only the integer index of retrieved entry is stored in a binary file in a custom SpectroVQ format (. vqms2). As the size of the codebook is known, each integer can be stored in log_2_ bits. The quantizers are ranked hierarchically, with the first one capturing the most information, the user can specify how many quantizers they use, counting from the first one, to reconstruct the denoised spectrum. Using more quantizers leads to a higher reconstruction fidelity, but with a cost in storage size. In case the number of peaks in the denoised spectrum is fewer than 50% of that of the original spectrum, to encourage model to retain more information (e.g., peaks from a second peptide in a chimeric spectrum), we apply a second layer of compression. The residual peaks between the denoised and the original spectrum are encoded with the same model, thus converting it to another list of indices. We notice that using this improves peptide identifications when searching with SpectraST^15^ (Supplementary Figure S12). The metadata in the original mzML file is preserved in a separate parquet file. Tokens indexes of all spectra including the residual spectra are stored in the. vqms2 file. The spectrum indexes of the residual spectra are stored in a separate .pkl file, so a total of 3 files is produced during compression. To retrieve the spectrum, the whole procedure is reversed. The latent embedding is retrieved from the trained codebook by referring to the stored indices. Both the denoised spectrum and the residual spectrum (if any) are reconstructed based on the stored indices. The metadata is stored separately in a parquet file which can be used to reconstruct the original data file if needed.

### Datasets

We use the both the human HCD and the human HLA consensus library from PeptideAtlas^40^ as our dataset to train SpectroVQ. In total, the dataset contains 6.3 million spectra, each annotated as a different peptide with different charge state. 8% of the 6.3M spectra are non-tryptic peptides from immunopeptidomics experiments. The dataset is partitioned into both training and testing dataset (95:5). The consensus library contains spectra from peptides of length ranging from 10 to 45 amino acids, charge states ranging from 1 to 5, and precursor m/z, ranging from 250 m/z to 1800 m/z (Figure 1a). Shorter peptides with length less than 19 and charge states 2+ and 3+ comprise most of the dataset (64% from short peptides and 79% from charge states 2+ and 3+).

We use a total of three external evaluation datasets in this work. First, the nine-species dataset (https://zenodo.org/records/13685813, Version v2) was used to evaluate SpectroVQ’s denoising performance in unseen datasets. As the training data only contains spectrum from human samples, evaluation using spectrum from other species can demonstrate its generalization ability. Second, a large dataset from the Human Proteome Project (PXD000561) was used to evaluation peptide information retain in embedding space of SpectroVQ. For this dataset, Spectroscape^20^ was used to construct a spectral archive, leading to clusters of spectra organized by spectral similarity and annotated by MSFragger^28^. This data structure enables us to compare massive number of spectra from identical peptides quickly. Third, the 3-species datasets are extracted from Bart et. al.^17^ (PXD028735). Three replicate runs of *E. coli*., human and yeast samples acquired on Orbitrap and TripleTOF 6600 mass spectrometers are used in this work as a real-life dataset to test SpectroVQ’s compression. This dataset is chosen because the dataset contains samples from 3 different organisms and its newer (2022) than our consensus library (2019). It ensures no information leakage into the training data, thus allowing a fair evaluation.

### Evaluation Metrics

Reconstruction fidelity is evaluated using the cosine similarity as the metric. We compute the cosine similarity between the square-rooted intensities of the original binned spectrum and the corresponding reconstructed binned spectrum. To evaluate denoising effectiveness, we define the fraction of canonical ion peaks as the ratio between the number of b and y ions and the total number of peaks in the spectrum. To avoid train/test leakage, the denoising evaluation presented are only the testing portion of the 6.3M consensus spectrum data. Silhouette and the Davies-Bouldin score are calculated using the scikit-learn python library. Token similarity is evaluated by calculating the portion of identical tokens between two spectra. Jensen-Shannon (JS) divergence is calculated using the scipy python library. Token density from each peptide group is first calculated with the normalized occurrence of each token. JS divergence between token densities from peptide group pairs are then calculated. The intersection of theoretical b,y fragments from two different peptides is treated as the number of overlapping canonical fragments. Annotation of the peaks to enable canonical ion fraction calculation (Figure 3) is performed using Pyteomics^39^. For simplicity, only b,y fragments with charge state up to 2+ is counted. Isotopic peaks are not considered. No Neural losses are considered. Meanwhile, peak annotation beyond b,y ions (Figure 3a, 4d) are annotated using Quetzal^41^. Ion peaks annotated by Quetzal shares the same constrains as those from pyteomics^39^, but only allowing less than 2 neural losses to compensate for the decrease in mass precision after reconstruction. Due to the slow speed in for Quetzal web annotator, for Figure 4d, 20 spectra are sampled at each peptide length, and then passed to Quetzal for annotation. Unannotated ions (Figure 4d) are defined as peaks without any quetzal annotation.

### Spectral Library Searching

Spectral library searching was performed on the original and reconstructed dataset to evaluate the congruence of peptide and protein identifications between the two datasets. We used Seq2MS^24^ to predict reference spectra of all tryptic peptides in the proteomes of their respective organisms. *E. coli* (UP000000625), human (UP000005640) and yeast (UP000002311) protein databases were downloaded from UniProt (https://www.uniprot.org/). Only reviewed protein sequences were included. Decoy reversed sequences were added to databases using Fragpipe^28^ with 1:1 ratio. Protein digestion was performed *in silico* using protein digestion stimulator. (https://pnnl-comp-mass-spec.github.io/Protein-Digestion-Simulator/). We used SpectraST^15^ for spectral library searching and Comet^25^ for database searching. Cysteine carbamidomethylation was considered as a static modification and methionine oxidation was allowed as a variable modification. Fragment bin size was set to 0.1 Th to match the highest m/z resolution outputted by SpectroVQ. PepXML results were merged and validated using PeptideProphet and InterProphet in the Trans-Proteomic Pipeline^41^ (version 7.1.0).

### Searching spectra of different denoising levels

Instead of specifying a fixed number of quantizers to use for SpectroVQ compression, we varied the number of quantizers from 1 to 12 to generate 12 different versions of each spectrum with different levels of denoising. All denoised spectra with different levels of denoising were then searched using SpectraST, and InterProphet^19^ was employed to merge and validate the results, with the “number of sibling peptides” and “number of replicate spectra” models disabled.

## Data availability

The curated Peptide Atlas library and the Seq2MS-predicted library of the *E. coli*, yeast and human proteomes, and the trained SpectroVQ model weights are available at https://zenodo.org/records/20043624. The nine-species dataset are available from https://zenodo.org/records/13685813. The three species DDA datasets are available at https://www.ebi.ac.uk/pride/archive/projects/PXD028735 <u>with ProteomeXchange ID</u> <u>PXD028735</u>. Reference proteome database for the three species *Homo sapiens*, *E. coli,* and *S. cerevisiae* are available on UniProt https://www.uniprot.org/proteomes with identifiers UP000005640, UP000000625 and UP000002311 respectively.

## Code availability

SpectroVQ’s code is available at https://github.com/kenll99minecart/SpectroVQ.

## Acknowledgements

This work is supported financially by the Research Grants Council of the Hong Kong Special Administrative Region Government (Grant Nos. 16307923, C5026-24G).

## Author contributions

J. H.W. L. was involved in the idea generation, data analysis, writing of the manuscript and implementation of SpectroVQ. A. H. was involved in preparation of the dataset using Spectroscape and provide advice on the algorithm development. H.L. conceived and supervised the project and guided the algorithm development and data presentation.

## Notes

### Competing Interest Statement

The authors have declared no competing interest.

## References

1. Guo, T., Steen, J. A. & Mann, M. Mass-spectrometry-based proteomics: from single cells to clinical applications. Nature 638, 901–911 (2025).

2. Deutsch, E. W. et al. The ProteomeXchange consortium in 2026: making proteomics data FAIR. Nucleic Acids Res. 54, D459–D469 (2026).

3. Perez-Riverol, Y. et al. The PRIDE database at 20 years: 2025 update. Nucleic Acids Res. 53, D543–D553 (2025).

4. Martens, L. et al. mzML—a Community Standard for Mass Spectrometry Data. Mol. Cell. Proteomics MCP 10, R110.000133 (2011).

5. Yang, R., Chen, X. & Ochoa, I. MassComp, a lossless compressor for mass spectrometry data. BMC Bioinformatics 20, 368 (2019).

6. Hanau, F., Röst, H. & Ochoa, I. mspack: efficient lossless and lossy mass spectrometry data compression. Bioinformatics 37, 3923–3925 (2021).

7. Bhamber, R. S., Jankevics, A., Deutsch, E. W., Jones, A. R. & Dowsey, A. W. mzMLb: A Future-Proof Raw Mass Spectrometry Data Format Based on Standards-Compliant mzML and Optimized for Speed and Storage Requirements. J. Proteome Res. 20, 172–183 (2021).

8. Van Den Bossche, T., et al. mzPeak: Designing a Scalable, Interoperable, and Future-Ready Mass Spectrometry Data Format. J. Proteome Res. 24, 5329–5335 (2025).

9. Du, P. et al. A noise model for mass spectrometry based proteomics. Bioinformatics 24, 1070–1077 (2008).

10. Awan, M. G. & Saeed, F. MS-REDUCE: an ultrafast technique for reduction of big mass spectrometry data for high-throughput processing. Bioinformatics 32, 1518–1526 (2016).

11. Seneviratne, A. J. et al. Improved identification and quantification of peptides in mass spectrometry data via chemical and random additive noise elimination (CRANE). Bioinformatics 37, 4719–4726 (2021).

12. Kong, F. et al. Denoising Search doubles the number of metabolite and exposome annotations in human plasma using an Orbitrap Astral mass spectrometer. Nat. Methods 22, 1008–1016 (2025).

13. Banerjee, S., Nakrani, P., Singh, A. & Wangikar, P. P. DuReS: An R Package for Denoising Experimental Tandem Mass Spectra and Metabolite Annotation. Anal. Chem. 97, 11986–11992 (2025).

14. Zeghidour, N., Luebs, A., Omran, A., Skoglund, J. & Tagliasacchi, M. SoundStream: An End-to-End Neural Audio Codec. Preprint at 10.48550/arXiv.2107.03312 (2021).

15. Lam, H. et al. Building Consensus Spectral Libraries for Peptide Identification in Proteomics. Nat. Methods 5, 873–875 (2008).

16. Wen, B. & Noble, W. S. A multi-species benchmark for training and validating mass spectrometry proteomics machine learning models. Sci. Data 11, 1207 (2024).

17. Van Puyvelde, B. et al. A comprehensive LFQ benchmark dataset on modern day acquisition strategies in proteomics. Sci. Data 9, 126 (2022).

18. Le Roux, J. & Sade, J. arcMS: transformation of multi-dimensional high-resolution mass spectrometry data to columnar format for compact storage and fast access. Bioinforma. Adv. 4, vbae160 (2024).

19. Shteynberg, D. et al. iProphet: Multi-level Integrative Analysis of Shotgun Proteomic Data Improves Peptide and Protein Identification Rates and Error Estimates. Mol. Cell. Proteomics MCP 10, M111.007690 (2011).

20. Wu, L., Hoque, A. & Lam, H. Spectroscape enables real-time query and visualization of a spectral archive in proteomics. Nat. Commun. 14, 6267 (2023).

21. Bittremieux, W., May, D. H., Bilmes, J. & Noble, W. S. A learned embedding for efficient joint analysis of millions of mass spectra. Nat. Methods 19, 675–678 (2022).

22. Litsa, E. E., Chenthamarakshan, V., Das, P. & Kavraki, L. E. An end-to-end deep learning framework for translating mass spectra to de-novo molecules. Commun. Chem. 6, 132 (2023).

23. Liu, K., Ye, Y., Li, S. & Tang, H. Accurate de novo peptide sequencing using fully convolutional neural networks. Nat. Commun. 14, 7974 (2023).

24. Chan, C. M. J. & Lam, H. Merging Full-Spectrum and Fragment Ion Intensity Predictions from Deep Learning for High-Quality Spectral Libraries. J. Proteome Res. 22, 3692–3702 (2023).

25. Eng, J. K. et al. A Deeper Look into Comet—Implementation and Features. J. Am. Soc. Mass Spectrom. 26, 1865–1874 (2015).

26. Su, D., et al. Token Assorted: Mixing Latent and Text Tokens for Improved Language Model Reasoning. Preprint at 10.48550/arXiv.2502.03275 (2025).

27. Li, J., et al. Discrete Tokenization for Multimodal LLMs: A Comprehensive Survey. Preprint at 10.48550/arXiv.2507.22920 (2025).

28. Kong, A. T., Leprevost, F. V., Avtonomov, D. M., Mellacheruvu, D. & Nesvizhskii, A. I. MSFragger: ultrafast and comprehensive peptide identification in mass spectrometry–based proteomics. Nat. Methods 14, 513–520 (2017).

29. Goldman, S., Li, J. & Coley, C. W. Generating Molecular Fragmentation Graphs with Autoregressive Neural Networks. Preprint at 10.48550/arXiv.2304.13136 (2024).

30. Défossez, A., Copet, J., Synnaeve, G. & Adi, Y. High Fidelity Neural Audio Compression. Preprint at 10.48550/arXiv.2210.13438 (2022).

31. Tagliasacchi, M., Li, Y., Misiunas, K. & Roblek, D. SEANet: A Multi-modal Speech Enhancement Network. Preprint at 10.48550/arXiv.2009.02095 (2020).

32. Défossez, A., et al. Moshi: a speech-text foundation model for real-time dialogue. Preprint at 10.48550/arXiv.2410.00037 (2024).

33. Touvron, H., et al. Llama 2: Open Foundation and Fine-Tuned Chat Models. arXiv.org https://arxiv.org/abs/2307.09288v2 (2023).

34. Akiba, T., Sano, S., Yanase, T., Ohta, T. & Koyama, M. Optuna: A Next-generation Hyperparameter Optimization Framework. arXiv.org https://arxiv.org/abs/1907.10902v1 (2019).

35. Kong, F. et al. Denoising Search doubles the number of metabolite and exposome annotations in human plasma using an Orbitrap Astral mass spectrometer. Nat. Methods 22, 1008–1016 (2025).

36. Loshchilov, I. & Hutter, F. Decoupled Weight Decay Regularization. Preprint at 10.48550/arXiv.1711.05101 (2019).

37. Xiong, R., et al. On Layer Normalization in the Transformer Architecture. Preprint at 10.48550/arXiv.2002.04745 (2020).

38. Li, Y. et al. Spectral entropy outperforms MS/MS dot product similarity for small-molecule compound identification. Nat. Methods 18, 1524–1531 (2021).

39. Levitsky, L. I., Klein, J. A., Ivanov, M. V. & Gorshkov, M. V. Pyteomics 4.0: Five Years of Development of a Python Proteomics Framework. J. Proteome Res. 18, 709–714 (2019).

40. Deutsch, E. W. The PeptideAtlas Project. Methods Mol. Biol. Clifton NJ 604, 285–296 (2010).

41. Deutsch, E. W. et al. Trans-Proteomic Pipeline: Robust Mass Spectrometry-Based Proteomics Data Analysis Suite. J. Proteome Res. 22, 615–624 (2023).

