## Supplementary Figures for "SpectroVQ: Noise-Aware Compression of Proteomics Data via Vector-Quantized Deep Learning improves MS/MS data storage and Peptide Identification"

*Correspondence to:

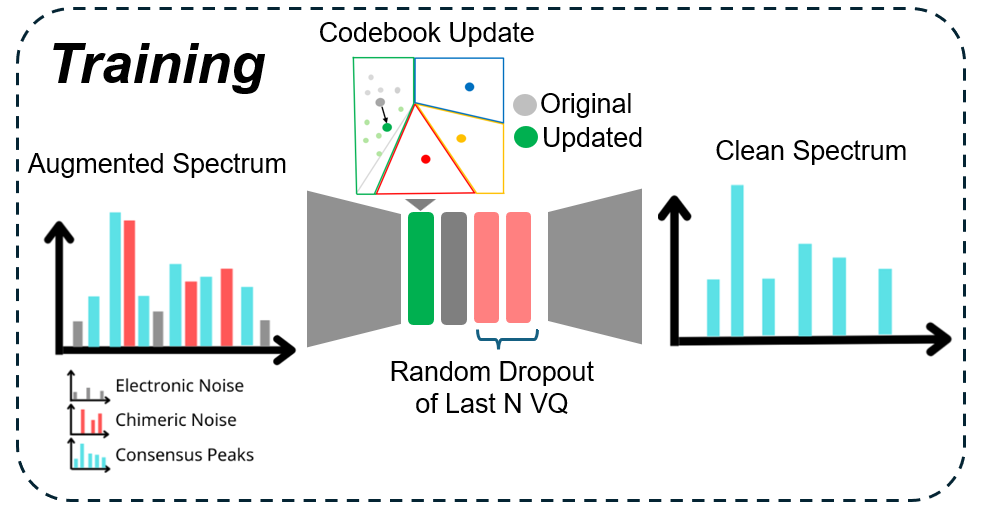
 **Figure S1 Training Scheme for SpectroVQ.** SpectroVQ is trained with a noise-to clean approach. Each spectrum has a chance to be augmented with electronic and chimeric peptide noise. During training, each codebook is updated with the trained latent, while the last N-quantizers are dropped out randomly to encourage the model to capture more information in the earlier quantizers, thereby achieving hierarchical compression.

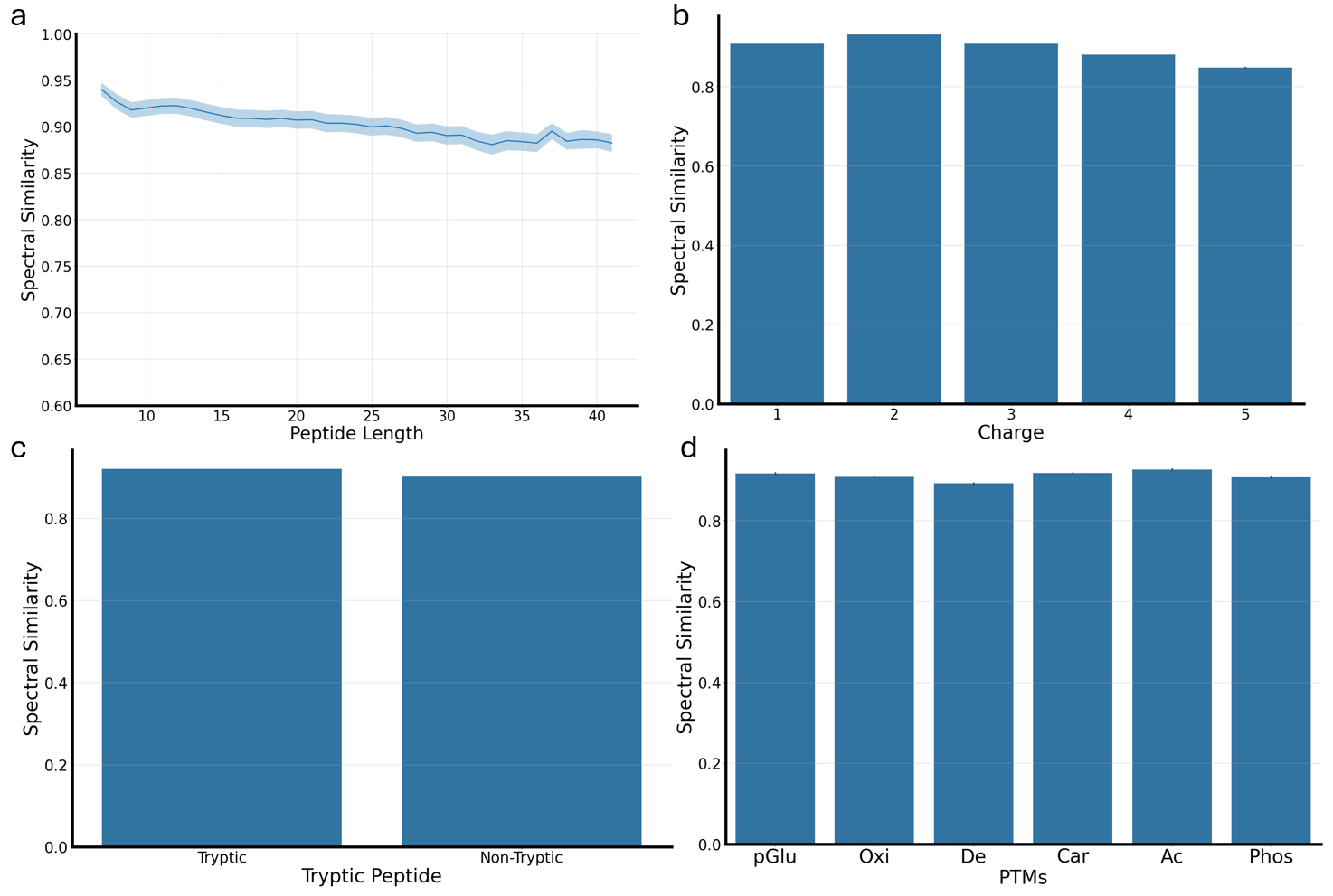

**Figure S2 Reconstruction Performance using SpectroVQ on the testing portion of the PeptideAtlas dataset.** Spectral similarity against spectrum from different peptide lengths **(A)**, charge state **(B),** tryptic and non-tryptic peptides portions **(C)** and Post-translational modifications **(D)** using 4 quantizers**.** Similar to the performance in Figure 2, excellent reconstruction fidelity was achieved for the vast majority of spectra (over 0.85 cosine similarity based on square root-transformed intensities). **Abbreviations**: Acetylation (Ac), Pyroglutamic acid modification (eGlu), Cysteine Carbamidomethylation (Car), Phosphorylation (Phos), Asparagine and Glutamine Deamidation (De), Methonine Oxidation (Ox)

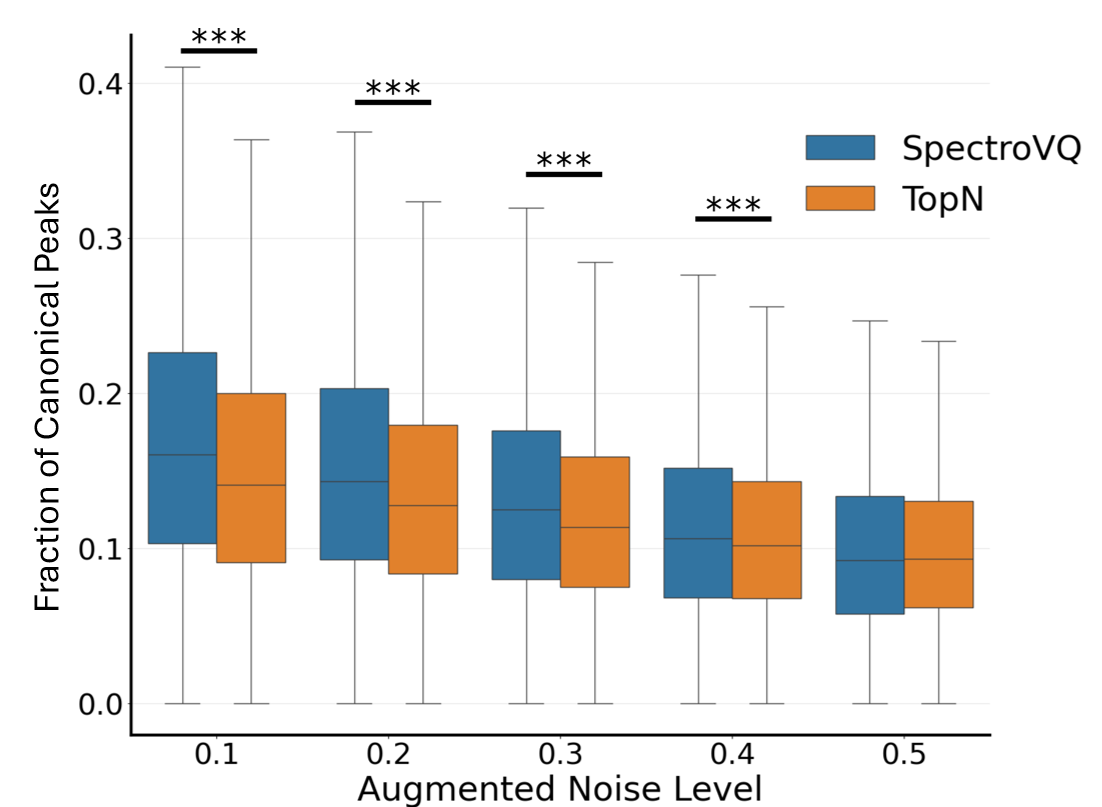
**Figure S3 The fraction of canonical peaks in the denoised spectra, versus the level of augmented noise for the testing portion of the PeptideAtlas dataset**. In a head-to-head comparison between SpectroVQ and a baseline denoising method of retaining the top N peaks (with N chosen to match the number of peaks retained by SpectroVQ), the fraction of canonical peaks retained by SpectroVQ is consistently higher across all noise level, though the advantage decreases as the noise level increases. This shows that SpectroVQ learned to retain the less intense canonical peaks over the more intense noise, whereas the Top N filter keeps the same number of peaks based on intensity alone. ****P* < 0.01 by two-tailed t-test.

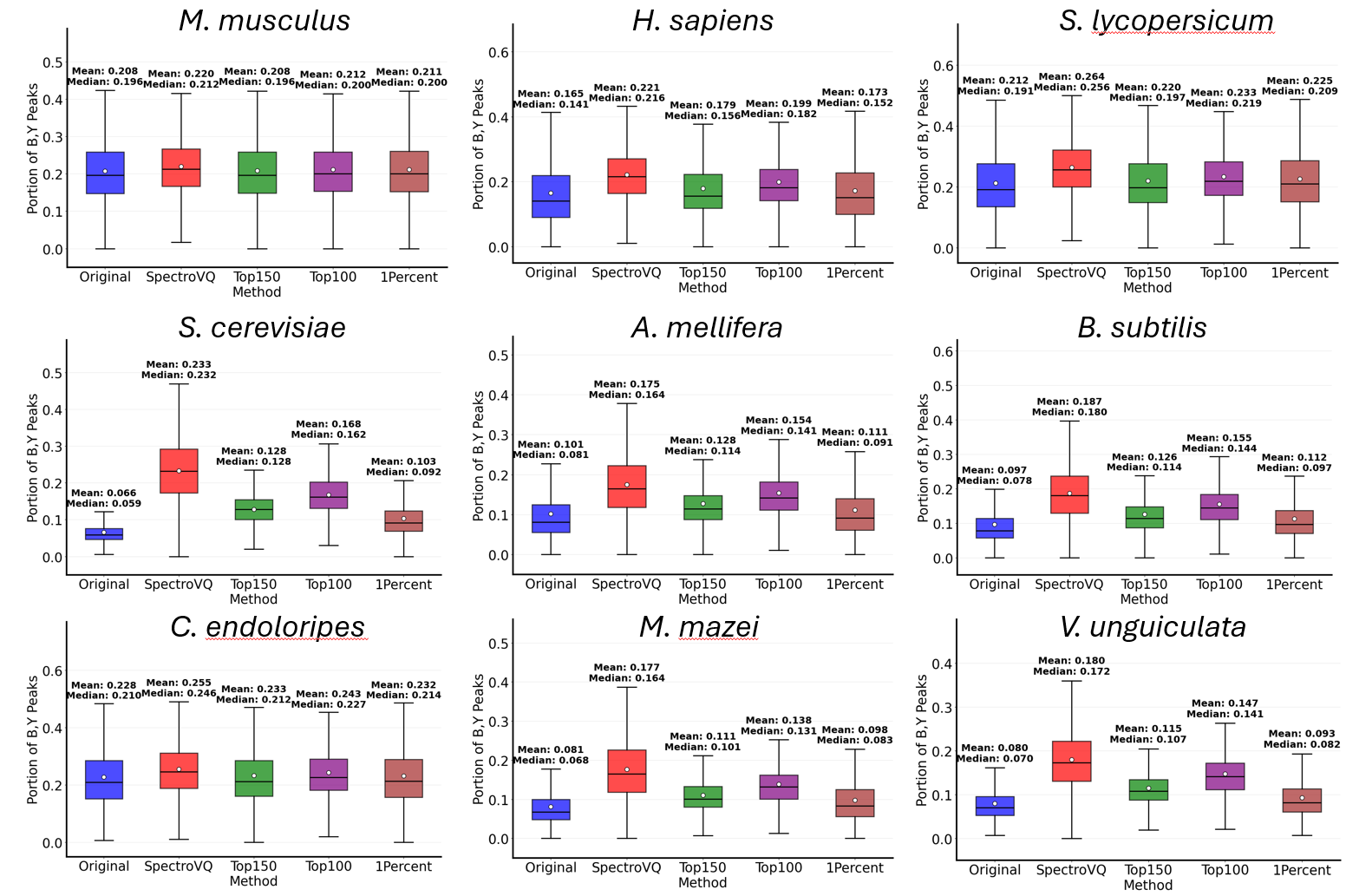

**Figure S4 Fraction of canonical peaks in the denoised spectra by different denoising strategies, for the nine-species dataset.** In terms of the ability to retain the canonical peaks, SpectroVQ consistently outrank other denoising methods of picking Top 150, Top 100 and thresholding peaks at 1% of the base peak intensity.

**Abbreviations:** *Vigna unguiculata* (*V. unguiculata*), *Methanosarcina mazei* (*M. mazei)*, *Candidatus endoloripes* (*C. endoloripes*), *Bacillus subtilis* (*B. subtilis)*, *Apis mellifera* (*A. mellifera.*), *Saccharomyces cerevisiae* (*S. cerevisiae*), *Mus musculus* (M. musculus), *Homo Sapiens* ( *H. sapiens*), *Salonum lycopersicum* (*S. lycopersicum*)

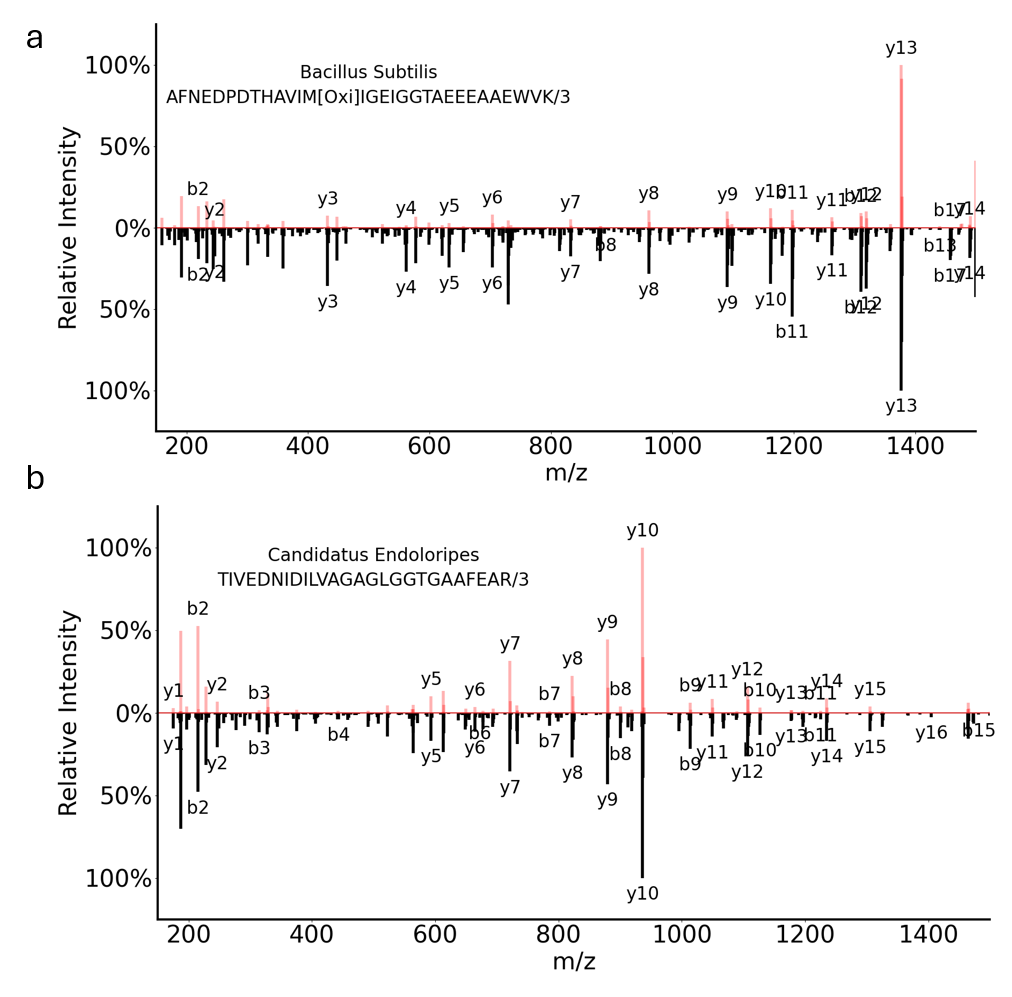

**Figure S5 Examples of denoised spectra in the nine-species datasets.** Denoised spectra by SpectroVQ (Red) and the corresponding original spectra (Black) in the *B. subtilis* **(A)** and *C. endoloripes* **(B)** are plotted against each other.

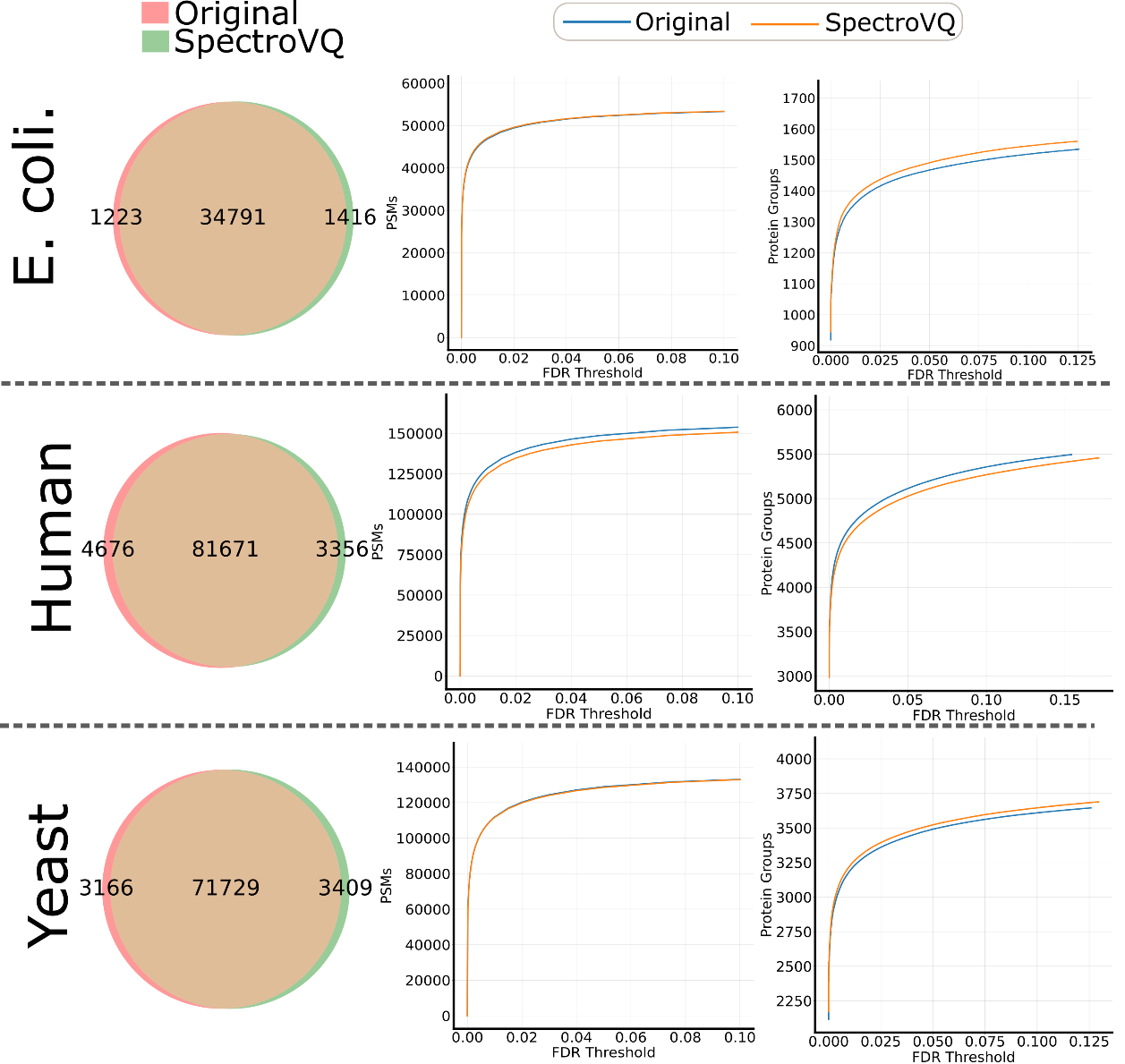

**Figure S6 Library search results of the original data versus SpectroVQ-processed data for the three-species dataset.** The number of spectra identified for the two sets of data (original and SpectroVQ) at 1% false discovery rate at PSM level shown as a Venn diagram, indicating >90% overlaps in PSMs (Left)**.** Number of PSMs (Middle) and protein groups (Right) versus FDR curves for the two sets of data show nearly identical identification rates between the two sets of data, indicating minimal biological information loss due to SpectroVQ compression.

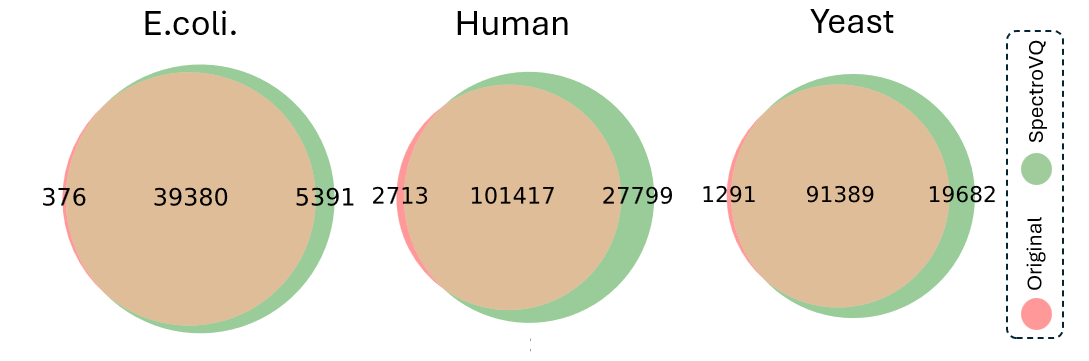

**Figure S7 Venn diagram comparing the SpectraST-identified spectra (at 1% false discovery rate at the PSM level) from the original spectra and from combining 12 denoised versions of the same spectra at different denoising level.** The multi-level searching strategy produces more than 15% more PSMs than searching the original data.

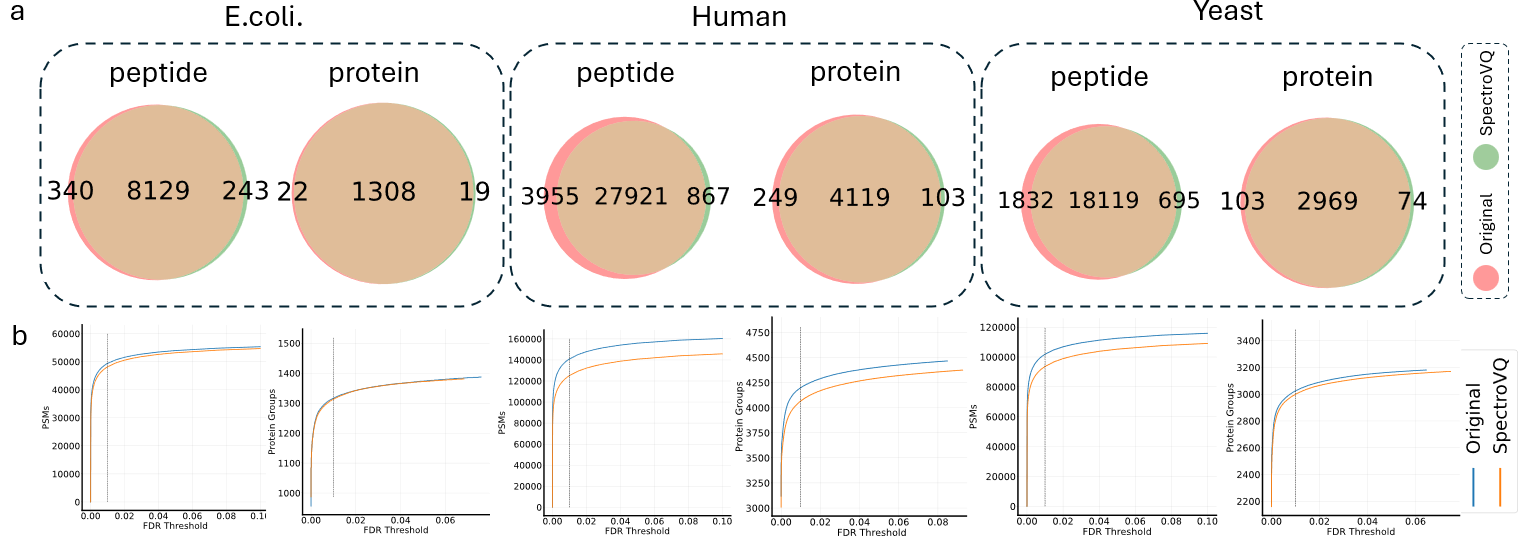

**Figure S8 Database search results of the original data and the corresponding SpectroVQ-processed data of the three-species dataset, using the Comet search engine** **(A)** Venn diagrams comparing the identified peptides and proteins between the original data and the SpectroVQ-processed data, at 0.01 false-discovery rate (FDR) **(B)** Number of identifications vs. FDR curves at the PSM (left) and protein levels.

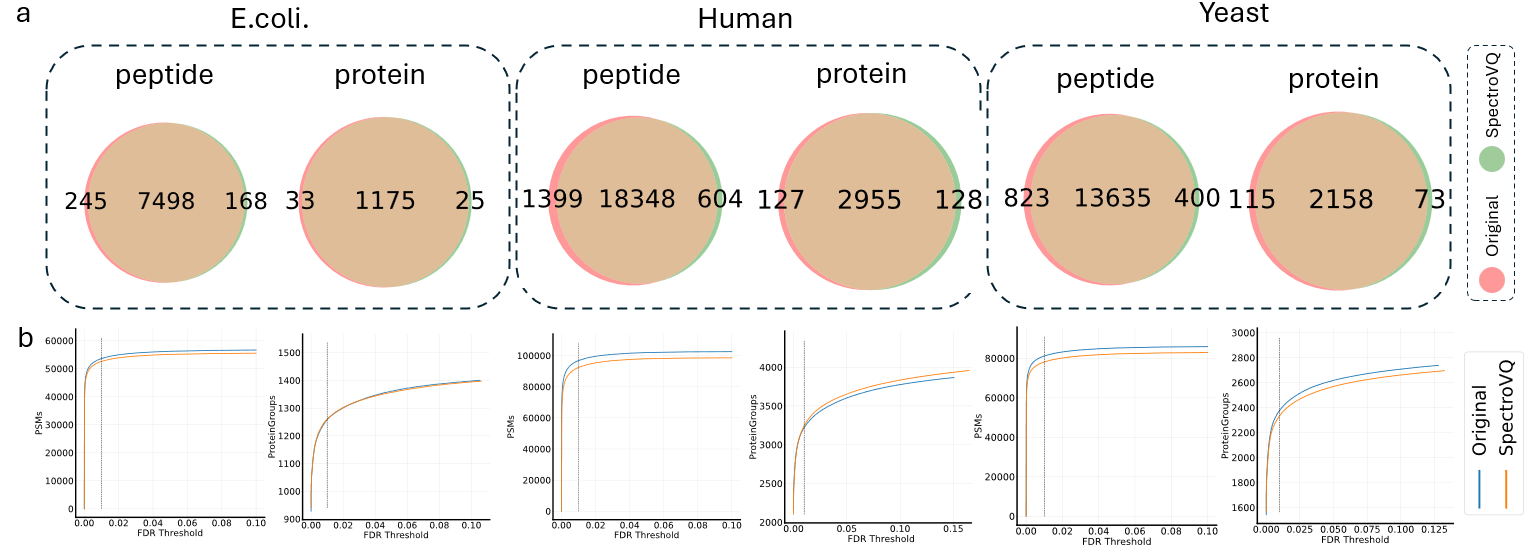
 **Figure S9 Database search results of the original data and the corresponding SpectroVQ-processed data of a TripleTOF 6600 dataset, using the Comet search engine (A)** Venn diagrams comparing the identified peptides and proteins between the original data and the SpectroVQ-processed data, at 0.01 false-discovery rate (FDR) **(B)** Number of identifications vs. FDR curves at the PSM (left) and protein levels.

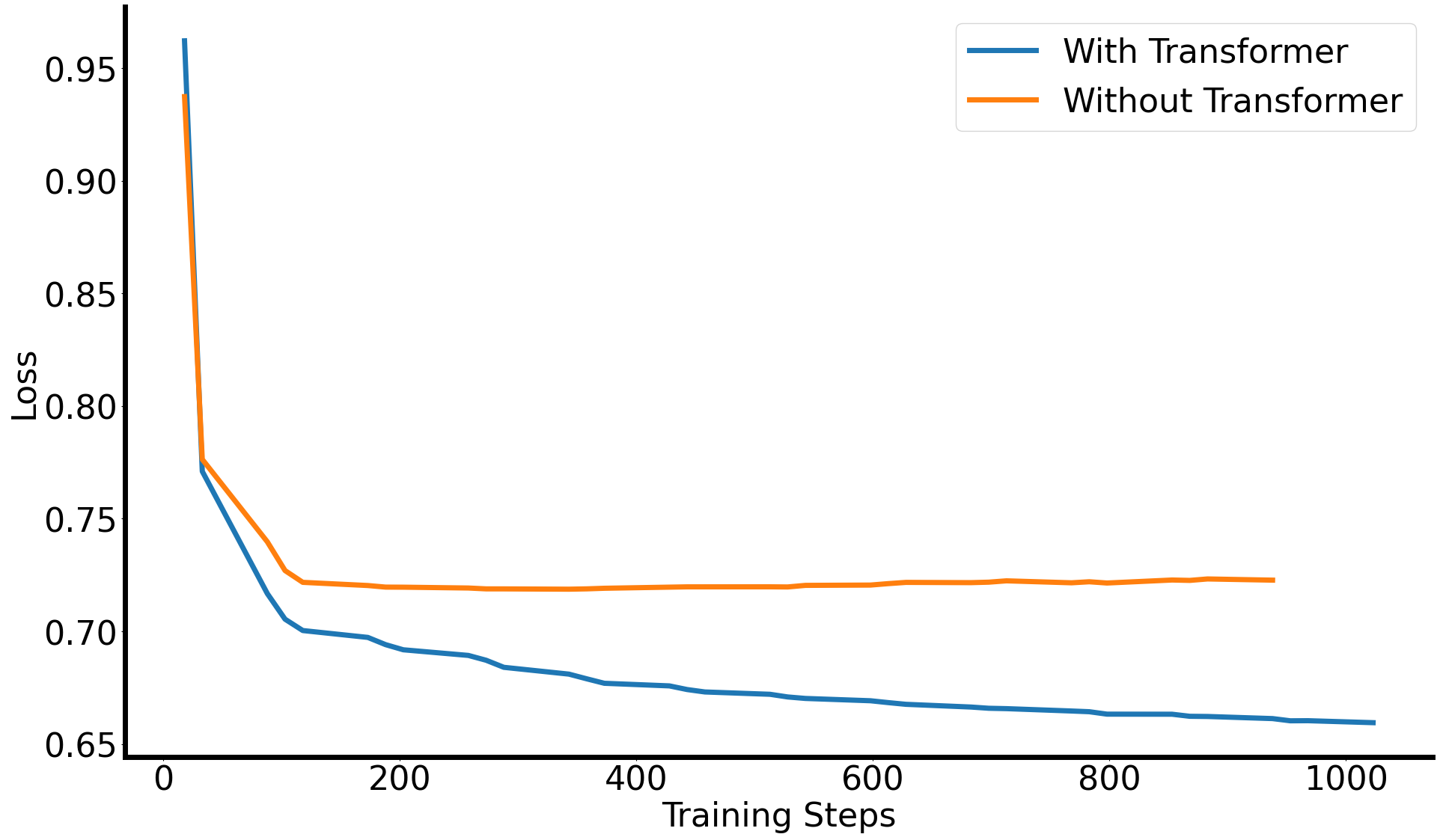

**Figure S10 Ablation study of SpectroVQ’s transformer layer.** Training epoch loss curves for SpectroVQ with and without the transformer layer show that adding the transformer layer greatly reduces training loss.

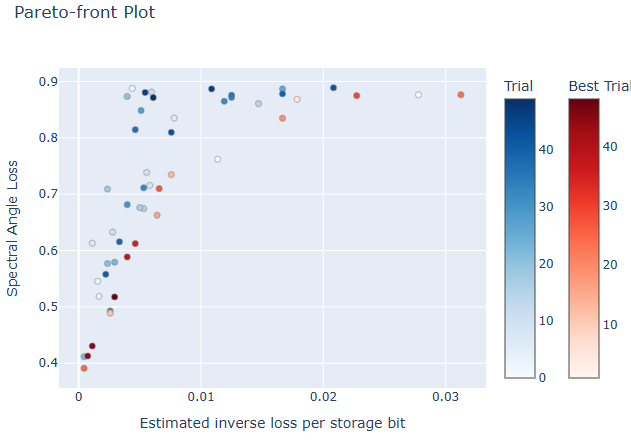

**Figure S11 Pareto front plot of the hyperparameter tuning process of SpectroVQ.** A trade-off between spectrum storage size (estimated inverse loss per storage bit) and reconstruction loss (spectral angle loss) can be observed. Optimized hyperparameters can be chosen from “best trial” settings located on the Pareto front.

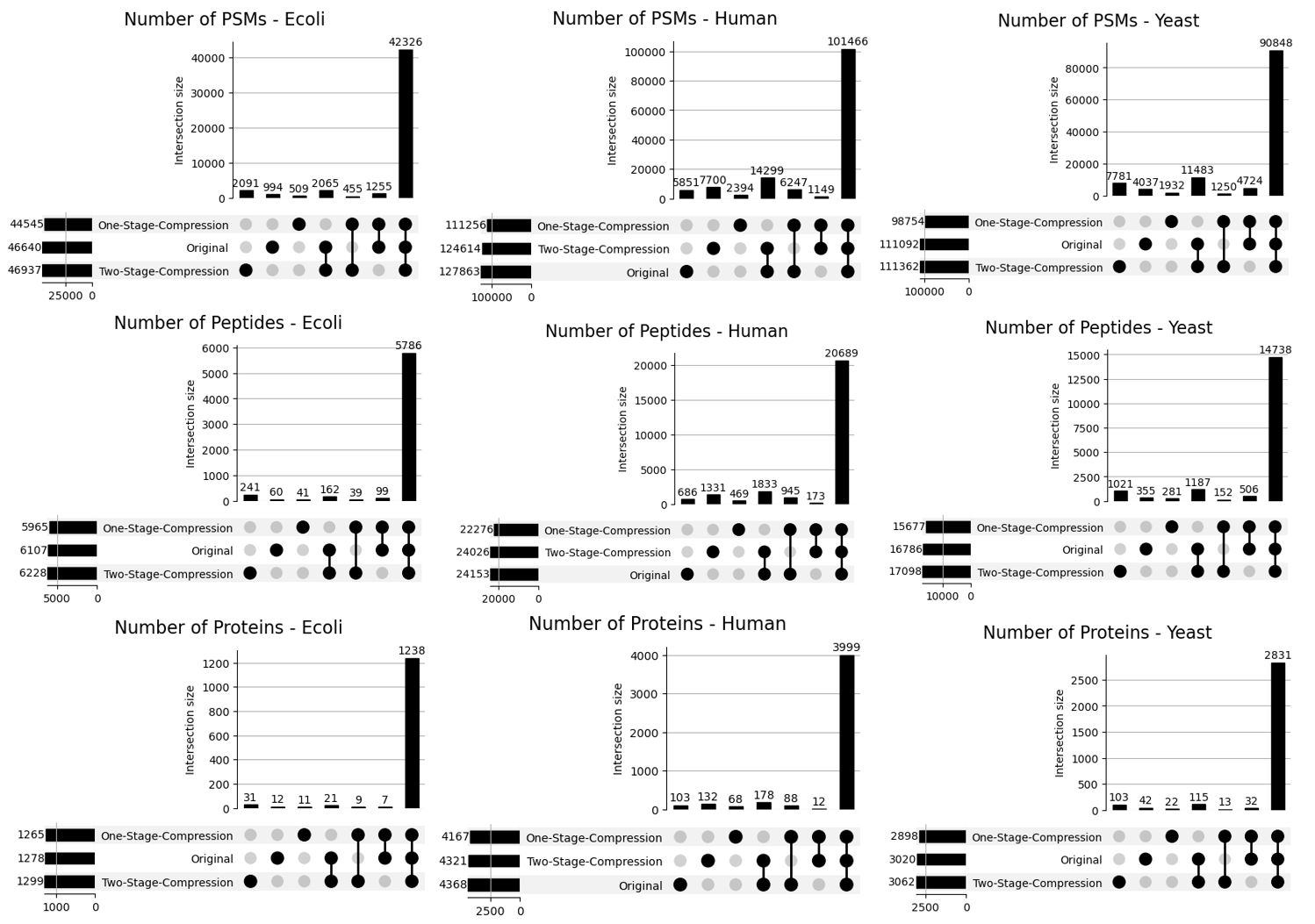

**Figure S12 Upsets plots for the identified Spectrum in Ecoli (Left), Human (Middle) and Yeast (Right) sample. The number of peptide-spectrum matches, peptides and proteins are filtered by 0.01 false discovery rate.** Compared to the original and simply using SpectroVQ’s model output, adding the residual spectrum to the raw denoised spectrum in SpectroVQ improves search results in all the 3-species dataset.

| **Hyperparameter** | **Value** |
| --- | --- |
| Total Number of Quantizers | 12 |
| Number of Channels (initial convolution layer) | 32 |
| Number of Residual Layers | 1 |
| Quantization Vector Length | 256 |
| Convolution kernel size | 6 |
| Stride Ratios | [9,7,4,2] |
| Codebook Size | 1024 |
| Activation Type | ELU |
| Final Activation Layer | Sigmoid |

**Table S1 Full list of hyperparameters of SpectroVQ**
